# Population-Level Genetic Variation Shapes Generative Brain Mechanisms

**DOI:** 10.1101/2023.08.31.555714

**Authors:** Alicja Monaghan, Danyal Akarca, Duncan E. Astle

**Author notes:** Corresponding authors: Alicja Monaghan, MRC Cognition and Brain Sciences Unit, 15 Chaucer Road, Cambridge, CB1 2HW.

## Abstract

The structural organisation of the human brain emerges probabilistically as we develop. Due to the inherent complexity of the human brain, understanding the forces that shape this probabilistic emergence remains one of the central challenges of systems theory and neuroscience. Across 2153 children (9-11 years old) we used a computational model to simulate the formation of structural brain connectivity, conceptualised as a trade-off between the cost of new connections η and their topological value γ. We then triangulated this population-level neuroimaging and computational modelling with genomics. For each participant we assessed their genetic propensity for cognitive ability by calculating polygenic scores. Modelled parameters differed systematically for participants depending upon their genetic propensity. Those with the highest genetic propensity had a significantly weaker η term – put simply, their networks emerged with a weaker distance penalty. Strikingly, this softer distance penalty produces more stochastic, diverse, and efficient networks. Furthermore, across the sample, overlapping biological and cellular pathways between polygenic scores and each child’s optimal η-γ trade-off emerged. This application of computational modelling demonstrates a converging genomic basis for structural brain development and cognitive ability across the population, providing a mechanistic explanation of how and why characteristic network topologies emerge from children at the extreme distributions of polygenic scores, and why they might predict cognitive ability.

## INTRODUCTION

Across development the human brain organises itself across multiple spatial and temporal scales, supporting an increasingly elaborate repertoire of cognitive and behavioural abilities. However, this process of organisation is highly variable, with this variability associated with substantial individual differences across several broad domains of cognition, including executive functioning, memory, language, and fluid reasoning (Jung & Haier, 2007; Siugzdaite et al., 2020). Indeed, many of the cognitive difficulties that cut across multiple neurodevelopmental populations vary in line with structural brain organisation (Rudie et al., 2013). But what fundamental mechanisms shape the emergence of brain organisation, and its variability, in the first place?

Brain structure describes the configuration and properties of white matter tracts and surrounding gray matter, comprised of myelinated axons and cell bodies, respectively. Across development this structure undergoes periods of dynamic change. Anatomical connections are pruned and refined (Huttenlocher & Dabholkar, 1997), with tract-specific and whole-brain trajectories varying by age, as the developing brain systematically organises itself. Whole-brain white matter (WM) gradually increases non-linearly before peaking in young adulthood (Bethlehem et al., 2022). However, the rate varies across the brain. Myelination of commissural and projection tracts are consolidated by late adolescence, whilst association tracts continue to mature until young adulthood (Lebel & Beaulieu, 2011). These non-linear and tract-specific trajectories mean that across development, there are dynamic and non-stationary relationships between brain structure and cognitive development (Simpson-Kent et al., 2020).

Rather than focussing on the structures of individual brain regions or tracts, a connectome provides a way of representing the organisation of *networks*, with each brain region represented as a node connected by edges (Bullmore & Sporns, 2009). Graph theory, established by Euler (1741), can be used to describe these networks mathematically, allowing us to understand the organisational principles of the human brain (see Bullmore & Sporns, 2009). The human brain has several organisational hallmarks consistent across scales, including lognormal distributions of firing rates and synaptic weights (see Buzsáki & Mizuseki, 2014), and therefore edge weights. Another hallmark is hubs, made up of typically long-range nodes whose connectivity exceeds chance (reviewed by Oldham & Fornito, 2019; van den Heuvel & Sporns, 2011). The result is a network that can be described as ‘small world’ (Watts & Strogatz, 1998), balancing the efficiency of the network with wiring costs (Kaiser & Hilgetag, 2004). Adult-like topological hallmarks are present early in development, such as small-world topology by 2 years of age (see Gilmore et al., 2018; Hagmann et al., 2010). Instead, most developmental changes are based on refining the relationships between hubs, such as increased segregation and average node strength, alongside reduced segregation (Hagmann et al., 2010).

Graph theory is a powerful way of capturing organisational properties of networks but does not explain the *origins* of those properties. Generative network models (GNMs) simulate how a complex network arises from a sparse ‘seed’ network, upon which connections between nodes are added probabilistically until the observed connectome is simulated. The model’s first instantiation added new connections in order to minimise wiring costs (Kaiser and Hilgetag, 2004), with subsequent extensions adding a second constraint (Vértes et al., 2012; Betzel et al., 2016) where nodes with similar properties wire together to maximise efficiency. Put simply, this class of model allows for the compression of complex topology to a relatively simple economic trade-off, which unfolds over time. This in turn allows one to explore how that economic trade-off might operate differently across different model systems and individuals.

The application of GNMs is in its relative infancy, but the main lessons are as follows: Structural brain development is driven by connections between nodes with similar properties. This is consistent across multiple spatial scales, species and developmental timepoints, including human cerebral organoids (Akarca et al., 2022), mice (Carozza et al., 2022), and non-invasive neuroimaging of child and adult brains (Akarca et al., 2021; Betzel et al., 2016). Variations in GNM specifications designed to reflect development and genomic patterning of the brain, such as wiring costs varying with head size and genetic constraints, have been shown to more accurately model structural brain organisation than static GNMs with fixed wiring costs and without genetic constraints (Arnatkeviciute et al., 2021; Oldham et al., 2022). Individual differences in η and γ parameter combinations have predictive validity across different cognitive domains, paradigms, and psychopathology: they differentially predict cognition in neurodevelopmentally-diverse children (Akarca et al., 2021), adaptive stress responses in early life adversity paradigms (Carozza et al., 2022), and are detuned in schizophrenia (Zhang et al., 2021).

One crucial influence on both our cognition and structural brain development is our genetics. Genetic influences on structural brain development continue throughout childhood and adolescence. Different aspects of cortical morphology are patterned by gene expression gradients, with this patterning systematically shifted in those with copy number variant disorders (Seidlitz et al., 2020). Indeed the genetic basis of structural brain development likely overlaps with that of cognitive ability. Several genome-wide association studies (GWAS) for genes associated with cognitive ability also contribute towards structural brain maturation (Savage et al., 2018; Sniekers et al., 2017).

One approach to assess the genetic basis for complex phenotypic traits, such as cognitive ability, is to use polygenic scores (PGSs). These require substantially smaller sample sizes than GWAS studies. A PGS measures an individual’s genetic predisposition towards an outcome by summing each single nucleotide polymorphism (SNP; changes in single DNA nucleotides within a gene), weighted by GWAS effect sizes (Choi et al., 2020).

The aim of the current study was to combine these ingredients to understand *how* common genetic variants linked to cognition shape generative brain mechanisms. We compared 5 GNMs (Betzel et al., 2016) to determine the best-fitting model for structural brain organisation, before fitting that model to each individual participant. Next, we used PGSs for cognitive ability to test whether and how common genetic variants linked to cognition change the generative parameters within the model. Finally, we tested the overlap between the functional role of the genes that contribute to those PGSs and regional gene expression data from the Allen Human Brain Atlas (AHBA; Seidlitz et al., 2020) using a comparative pathway enrichment analysis.

## RESULTS

### PGSs for Cognitive Ability Account for Up to 4% of Variance in g Factor Loadings

The first step was to create a PGS for cognitive ability. Taking our base dataset (Savage et al., 2018), we created PGSs (full details in Section 5.5) within a subset of the ABCD dataset who had passed genomic quality control (N = 6528, 52.88% male, 81.53% European), aged between 8.92 and 11.08 years old (Mean = 9.93 ± .62 years). As our phenotype we took the first principal component (PC) across seven different tasks from the NIH toolbox, which had been administered to all participants, spanning attention (Eriksen & Eriksen, 1974; Rueda et al., 2004; Weintraub et al., 2013), set shifting (Zelazo, 2006), working memory (Tulsky et al., 2014), episodic memory (Dikmen et al., 2014), language (Gershon et al., 2013; Woollams et al., 2011), and processing speed (Carlozzi et al., 2015). Task details can be found in **Supplementary Table 1**. A 5-factor solution accounted for 85.04% of variance in the cognitive battery (full details in Section 5.11). The first PC – hereafter labelled ‘*g’*-captured 37.76% of variance in cognitive scores. Crucially, this variance is evenly distributed across all tasks (**Supplementary Table 2**), as we would expect with a *g* factor (see Deary et al., 2010). The PGSs were a significant positive predictor of PC1 loadings across both European (*t* = 14.38, *p* = 5.12 x 10^-46^, β = 13024.31, *R^2^* = .04) and non-European (*t* = 4.56, *p* = 5.58 x 10^-6^, β = 12194.43, *R^2^* = .02) subsets (**Supplementary Table 3**) with clumping inclusion thresholds of *p* = .10 and *p* = .20, respectively (**Supplementary Table 4**). These loadings are broadly in line with multiple other polygenic scores of cognitive and behavioural phenotypes (Genç et al., 2021; Lett et al., 2020; Loughnan et al., 2020; Savage et al., 2018; Sniekers et al., 2017) and indicates that we have accurately captured the established common genetic variants that are associated with cognitive ability in the wider population.

### Properties of Individual-Level Consensus-Thresholded Structural Connectomes

The second step was to generate and threshold group-level and individual-level structural connectomes for a subset of 2154 ABCD participants with high-quality T1w, genomic, resting-state functional MRI (rsfMRI) and tractography data (N = 2154, 52.09% male, 55.85% White) aged between 8.92 and 11.08 years old (Mean = 9.91 ± .62 years). We stratified the 4306 participants with high-quality T1w, genomic, rsfMRI and tractography data on cognitive ability, age, and sex (see **Supplementary Section 1**).

We conducted deterministic tractography for each participant (**Figure 1a**), and reconstructed structural connectomes (**Figure 1b**) as streamline counts (edges) between regions (nodes). Foreshadowing our later analysis, to maximise spatial coverage and minimise noise from the AHBA, we selected the Schaefer 100-node (17-network) parcellation. We applied a 60% consensus threshold to individual connectomes to remove spurious connections (de Reus & van den Heuvel, 2013), and applied a distance-dependent consensus threshold (Betzel et al., 2019), which maintains the distribution of shorter edges. This generated the group-level connectome which the GNMs (**Figure 1c**) simulated. We then thresholded individual connectomes at 27 streamlines to create individual-level connectomes which the GNMs attempted to simulate, with an average density of 6.55% (± .48%) across participants. This threshold was the same as previous GNM studies (Akarca et al., 2021; Betzel et al., 2016), and produced an average density comparable to a previous GNM study using a 108-node parcellation (Betzel et al., 2016). These thresholding steps were necessary because GNMs require binarized input matrices. The final thresholding step was a 95% consensus threshold to individual structural connectomes to generate a seed network for the group- and individual-level GNM analyses that we will describe later (full details in **Section 5.3**). Before computational modelling, we explored the characteristics of the empirical thresholded connectomes. Averaged across participants, high-degree (highly connected) regions included the left temporal pole, left prefrontal cortex and bilateral precuneus, whilst right somatosensory regions were less connected (**Figure 1d**). Long-range connectivity dominated the bilateral temporal poles, and short-range connectivity in the right somatosensory and motor regions (**Figure 1e**). Bilateral superior temporal lobules and temporal poles had the highest betweenness-centrality, and therefore appeared to mediate the greatest proportion of shortest paths between nodes (Newman, 2005), whilst right somatosensory and motor regions the least (**Figure 1f**). Regions with high clustering, or those which are fully connected to others within a community (Bullmore & Sporns, 2012), included the bilateral secondary somatosensory and motor S2 regions, whilst the least clustered included the right primary somatosensory and motor regions (**Figure 1g**). Descriptive statistics of local and global properties for the Brainnetome-246 and Schaefer-400 node parcellations are provided in **Supplementary Table 5**.

**Fig. 1.**
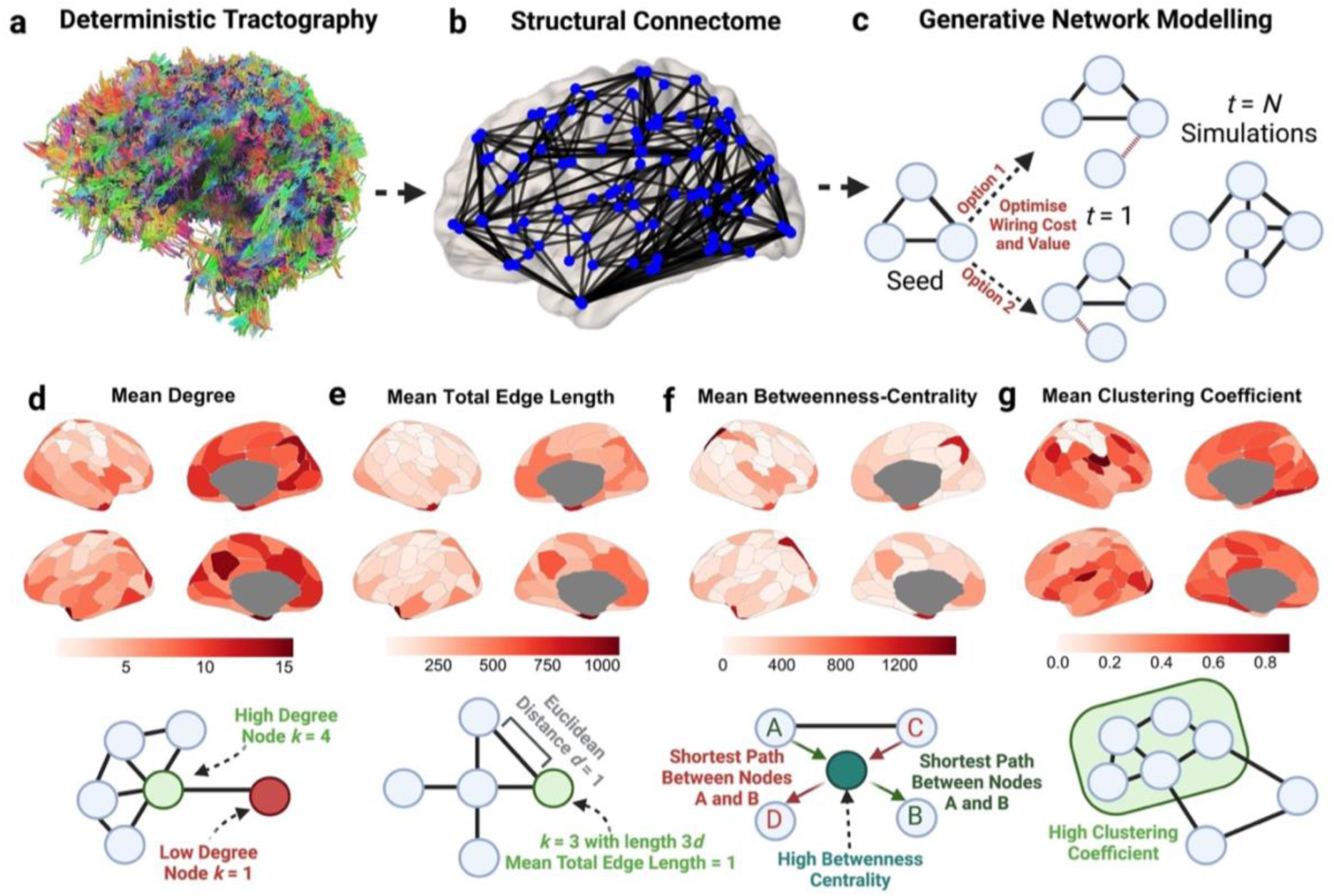
Empirical connectome formation and properties. **a** Using deterministic tractography on 2154 ABCD participants, we reconstructed individual structural connectomes as inter-regional streamline counts for the Schaefer 100-node parcellation. **b** These connectomes formed structural networks, made up of regions (nodes) connected to each other through edges. **c** Starting from a sparse seed network of connections common to all participants and thresholded at 27 streamlines per node, we simulated each participant’s structural connectome using generative network modelling. This technique adds one edge per iteration based on wiring probabilities, computed as a trade-off between the metabolic cost and network value of the new edge. After the first iteration (*t* = 1), wiring probabilities are updated, and single edges added until the simulated connectome had the same number (N) of edges as the observed empirical connectome. Averaged across all participants, the individual consensus-thresholded structural connectomes varied spatially in **d** mean degree (*k*, the number of connections each node has), **e** total edge length (the average Euclidean distance d of each edge connected to each node), **f** betweenness-centrality (the proportion of edges passing through a node that constitute a shortest path), and **g** clustering (nodes with similar properties tend to cluster together, forming neighbourhoods). Metrics **d-g** were used to evaluate simulation performance in subsequent analyses.

### Neighbour-Homophily Generative Network Models Provide the Best Account of Structural Brain Organisation Across 5 Models

The third step was to fit generative network models (GNMs) to the 2153 participants with high-quality thresholded structural connectomes. To enable comparisons between participants, all models used the same seed network of connections common to at least 95% of participants, with 1.58% density (see **Supplementary Figure 2**). Starting with a sparse seed network, GNMs simulate the formation of a network by probabilistically adding a single edge to unconnected nodes in the network, until the network has equal density to the observed connectome for that participant. At each iteration, a probability score is calculated that a new edge will be added between nodes i and *j*, as a trade-off between the wiring cost term η, which assesses the metabolic cost of adding a new edge, and a wiring value term γ, which measures the contribution of this new edge to the network topology. This is formalised as *P*(*i,j*) = *E*(*i,j*)^η^ x *K*(*i,j*)^γ^ (Betzel et al., 2016). *E*(*i,j*) represents the Euclidean distance between nodes *i* and *j*, reflecting how physical space constrains brain development and that longer-range connections incur greater metabolic costs (Bullmore & Sporns, 2012). Different ways of achieving economical brain organisation are implemented in GNMs through different wiring rules, represented by *K*(*i,j*) ^γ^, broadly grouped into 4 classes (**Figure 2a**): degree, clustering, homophily, and spatial. For example, in the case of two unconnected nodes *i* and *j*, the probability of a shared edge being added varies depending on the wiring rule, such as the number of edges each node has (degree), the proportion of their edges that are fully connected (clustering), similarity in connectivity profiles and shared neighbours (homophily), or simply their proximity (spatial). Therefore, each participant’s structural connectivity development can be summarised by a unique trade-off between wiring cost η and some topological value γ.

**Fig. 2.**
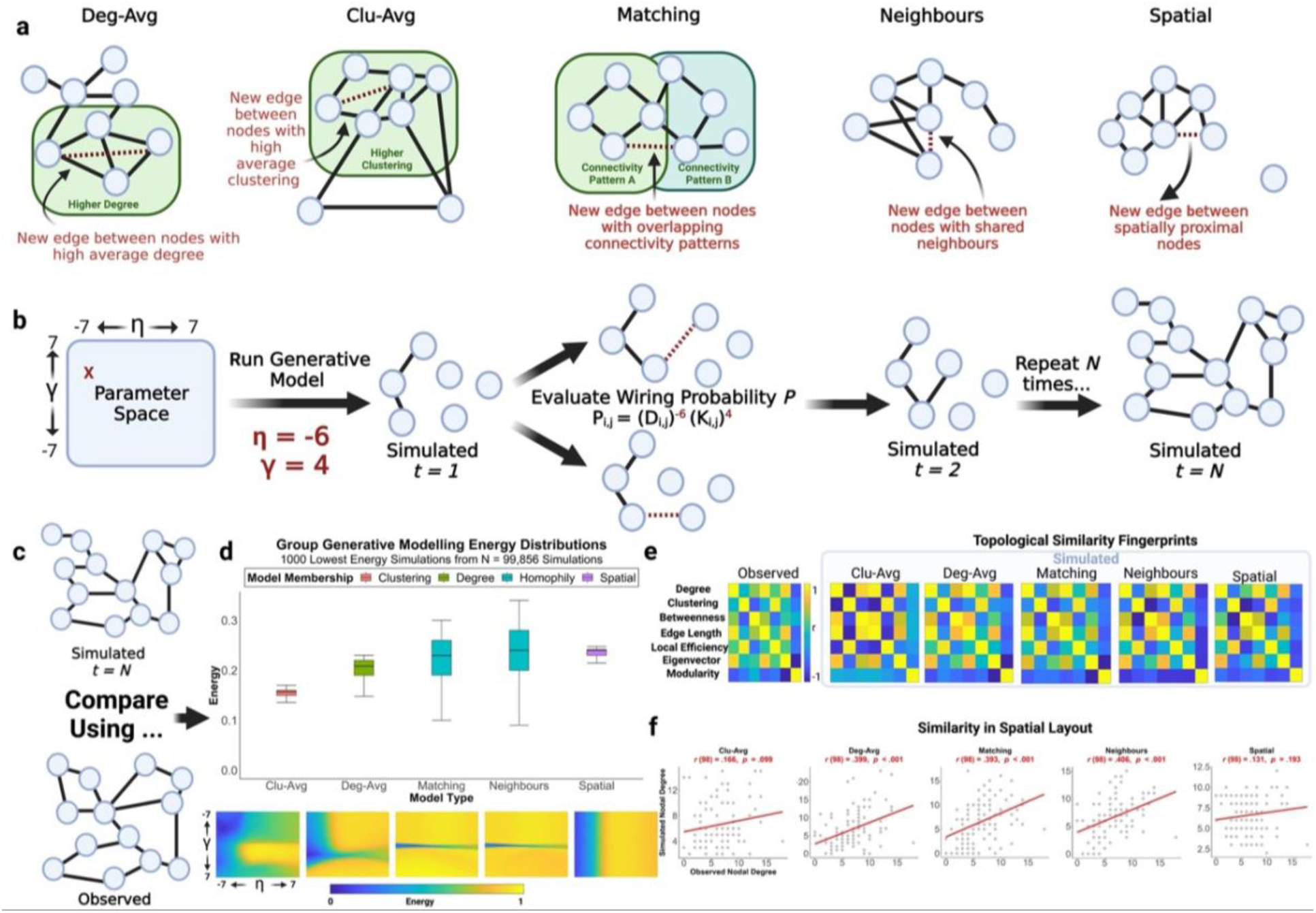
Generative network modelling protocol and evaluation. **a** 5 generative network modelling rules were evaluated. The deg-avg (degree average) model adds edges to nodes with high average degrees. The clu-avg (clustering average) model adds edges to nodes with high clustering coefficients. The homophily-matching model adds edges to nodes with a high matching index, or normalised similarity in connectivity profiles. The homophily-neighbour model adds edges to nodes sharing similar neighbours. The spatial model adds edges to proximal nodes. **b** A random wiring cost (η) and wiring value (γ) combination from a parameter grid search. This combination is used to run all 5 generative models, with N iterations. At each iteration, the wiring probability between each connected and unconnected node is computed. **c** The simulated connectome is compared with the observed connectome using 3 metrics. **d** The first is model energy (middle), equal to the largest Kolmogorov-Smirnov statistic testing differences in distributions of edge length, clustering, betweenness-centrality, and edge length. Distributions are shown for 5 models across 4 model classes. Corresponding energy landscapes are visualised. The homophily model energy landscapes show convergence on a small parameter window, despite having a large energy range, in contrast to the other models. **e** The second metric is topological similarity, which assesses the ability of the simulated connectome to recapitulate the local distribution of nodal properties in the observed connectome. The fingerprint plotted is representative of the overall statistical trends for each model across all parameter combinations and 1000 simulations of the lowest-energy parameter combinations. **f** The third metric is similarity in spatial layout, equal to the correlation between observed and simulated nodal degree. Simulated degree representative of the mean Pearson correlation between simulated and observed degree across 1000 simulations of each wiring rule’s lowest-energy parameter combination is plotted.

Fitting multiple generative models to 2153 participants would consume massive computational resources, so we instead performed our model selection on the group connectome (Betzel et al., 2019). Based on discrepancies in the literature surrounding the best-performing generative network models, we first ran 100,000 simulations for the top-performing model classes based on the prior literature (Akarca et al., 2021; Betzel et al., 2016). This yielded 99,856 unique η [-7 ≤ η ≤ 7] and γ [-7 ≤ γ ≤ 7] combinations (**Figure 2b**). We evaluated the similarity between the simulated and observed connectomes using three different measures of fit (**Figure 2c**). The first is energy (**Figure 2d**), equal to the largest dissimilarity between synthetic and observed connectomes in terms of one of four nodal metrics: degree, clustering, betweenness-centrality, and edge length distributions (Betzel et al., 2016). Kolmogorov-Smirnov (KS) statistics are computed for each nodal metric, and integrated, to test compare the distribution of each nodal metric in the observed connectome to that in the simulated connectome. The lower the energy, the better the model fit. The homophily-neighbours model achieved the lowest energy (.090), with η as −2.911 and γ as .244, respectively, followed closely by the homophily-matching model (.100). Across the best 25 simulations for each model, the two homophily models performed best [*F*(4,124) = 577.11, *p* = 2.48 x 10^-77^], with follow-up post-hoc tests confirming that these two models performed better than the other model types (all *p* < .001), but equivalent relative to each other (*p* = .69). Despite the homophily models having the smallest energy, they also displayed the largest range for the lowest-energy 1000 simulations of .250 and .200, respectively. This suggests that homophily can produce very low energy simulations but are most sensitive to parameter tuning. As shown in the corresponding energy landscapes (**Figure 2d**), the 10% lowest-energy simulations for, for instance, the homophily-neighbours model corresponded to a small region of the convex optimisation space, bounded by a narrow range of η [-5.489 ≤ η ≤ −1.800] and γ [.111 ≤ γ ≤ .333], respectively.

Energy, whilst the most common measure for fitting GNMs (Akarca et al., 2021; Betzel et al., 2019; X. Zhang et al., 2021), only tests for similarities in global nodal distributions. Therefore, we also used a second evaluative metric, termed *topological similarity* (**Figure 2e**). This measures the extent to which the simulated connectome captures the *local* distribution of nodal properties (Akarca et al., 2022; Carozza et al., 2022). This is calculated as the correlation between simulated and observed connectomes of 7 nodal metrics: degree, clustering, betweenness-centrality, edge length, local efficiency, centrality, and modularity. This produces a ‘topological fingerprint’ or unique local nodal distribution map for each model. The Euclidean norm of the difference in topological fingerprints between the observed connectome and each simulated model produces a single topological similarity value. Because of the probabilistic nature of the GNM, we repeated this process 1000 times for the lowest energy parameter combination for each model. The models differ significantly on their topological dissimilarity [*F*(4,4999) = 2774.22, *p* < .001]. Homophily-neighbours is the best performing numerically, although both homophily models do well and equivalently (*p* = .209), relative to both the spatial (*p* < .001) and cluster-average models (*p* < .001). The degree-average model also performs well on this metric, performing significantly better than the homophily-matching model (*p* = .004) and equivalently to the homophily-neighbours model (*p* = .643).

The third and final evaluative metric (**Figure 2f**) is similarity in spatial layout, equal to the correlation between the nodal degree of simulated and observed connectomes. The larger the correlation, the better the model fit. Again, we repeated this calculation 1000 times for the lowest-energy parameter combinations for each model. This measure differed significantly across the models [*F*(4,4999) = 3466.96, *p* < .001], with the homophily models performing better than all other classes (all *p* < .001). The homophily-neighbours variant achieved the highest spatial correlation (mean *r* = .42), which is significantly better than all other models, including the homophily-matching variant (all *p* < .001). In summary, the homophily models perform particularly well across all three metrics. We carried forward the homophily-neighbours model for our subsequent analyses because it achieved an equivalent energy, and slightly better topological dissimilarity and spatial correlation, relative to the other homophily variant. Although it is important to note that these two models are mathematically extremely similar, which would explain why they perform so similarly. Lowest-energy eta and gamma combinations, alongside mean topological dissimilarity and correlations between simulated and observed nodal degree across 1000 simulations for the lowest-energy parameter combinations in the Schaefer 100-node, Brainnetome 246-node, and Schaefer 400-node parcellations are presented in **Supplementary Tables 6 and 7**, respectively.

### A Negative Distance Penalty and Positive Topological Term Provides the Best-Fit for Individual-Level Homophily-Neighbours Models

The fourth step was to fit the homophily-neighbours model to individual participants. To narrow down the parameter window, we included used η and γ limits which bounded the top 10% lowest energy simulations from the consensus networks ([-7.000 ≤ η ≤ −.200] and [-.067 ≤ γ ≤ .600]). This yielded 74,529 unique combinations of η and γ, which were used to run simulations for 2153 participants. From the original sample of 2154, we removed one participant from the subsequent fitting procedure because of incorrect connectome construction.

As shown in **Figure 3a**, energy for the optimal simulations ranged between .050 and .140 (Mean = .071 ± .007). Distance penalties η ranged between −4.325 and −1.025 (Mean = −3.007 ± .336). These were all negative and larger in magnitude compared to the positive topological terms γ, which ranged between .071 and .323 (Mean = .215 ± .027). The small standard deviations for optimal η and γ confirmed that the parameter search was successful in a convex optimisation parameter window in the energy landscapes. In terms of nodal wiring cost (**Figure 3b**), ‘expensive’ areas included the bilateral cingulate posterior gyri, precuneus, and posterior cingulate cortices, whilst ‘cheap’ areas included the bilateral temporal pole and lateral ventral prefrontal cortices. In terms of nodal wiring value (**Figure 3c**), high-value regions tended to be in the prefrontal cortex, including bilateral medial posterior prefrontal, left dorsal prefrontal, and left medial prefrontal cortices, whilst lower-value regions tended to be in the parietal cortex, including retrosplenial regions, postcentral and intra-parietal sulci. The simulations recapitulated several features of the observed connectomes (**Figures 3d-k**), such as degree [*r*(98) = .561, *p* < .001] and edge-length [*r*(98) = .431, *p* < .001]. Crucially, they also recapitulated the emergence of local statistics excluded from the energy equation and model selection, such as local efficiency [*r*(98) = .291, *p* = .003] and eigenvector centrality [*r*(98) = .705, *p* < .001], However, the simulations struggled to accurately model the distributions of participation coefficients, betweenness-centrality, and clustering, typically under-stepping.

**Fig. 3.**
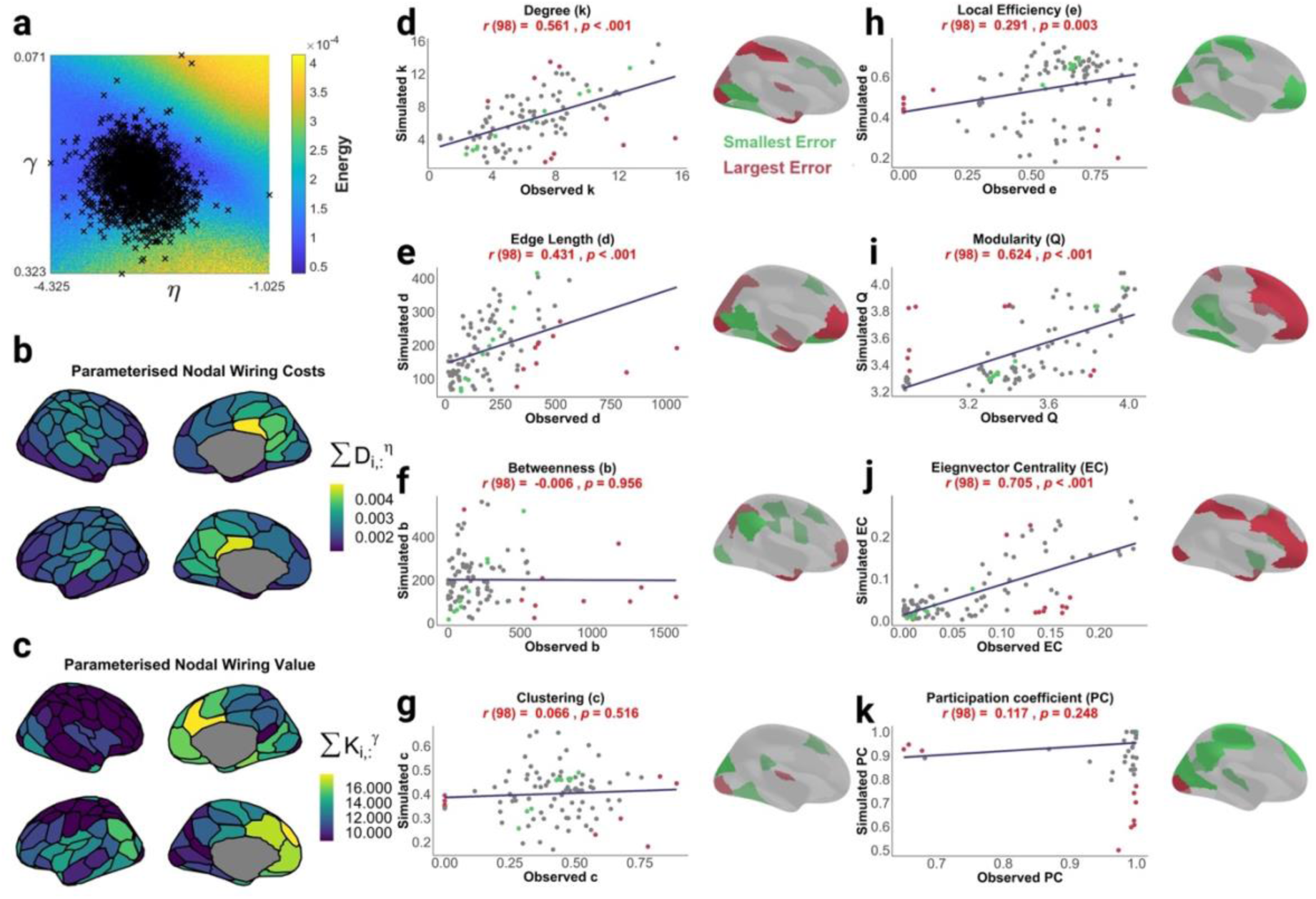
Lowest-energy individual homophily neighbour generative network models display regional heterogeneity in parameterised nodal wiring costs and value and recapitulate the distribution of local nodal statistics both included and excluded from the energy equation. **a** Lowest energy η and γ parameter combinations for 2153 participants, plotted against the associated energy landscape. **b** Averaged across participants, parameterised wiring costs for each node *i* are plotted as the sum of the Euclidean distance with all other nodes, parameterised by η. **c** Averaged across participants, parameterised wiring values for each node *i* are plotted as the normalised matching index with all other nodes, parameterised by γ. **d** Pearson correlations between simulated and observed nodal statistics, alongside regions with the smallest 10% (green) and largest 10% (red) absolute discrepancy between simulated and nodal statistics, are plotted for degree, **e** edge length, **f** betweenness-centrality, **g** clustering coefficient, **h** local efficiency, **i** modularity, **j** eigenvector centrality, and **k** participation coefficients. Statistics **d-g** were included in the energy equation, and the remainder were not.

### Do Common Genetic Variants Shape Generative Brain Mechanisms?

So far we have identified the best-fitting wiring rules and fit GNMs to individual participants. But how do generative mechanisms differ according to a participant’s PGS? A simple but powerful test of whether these common genetic variants shape the economic trade-off at the heart of the GNM is to compare parameters at the extremes. Therefore, we selected participants with the bottom 10%, and those with the top 10%, genetic propensity for cognitive ability, and compared the parameters for their best-fitting models (**Figure 4a**). Group membership (bottom versus top 10%) had a significant and selective impact on GNM parameters: η differed significantly [*t*(290) = −2.520, *p*= 0.012] between the groups, but γ did not [*t*(290) = −.194, *p*= 0.846]. Put simply, those with the highest genetic propensity had a significantly *softer* distance penalty, relative to their counterparts (**Figure 4b**).

**Fig 4.**
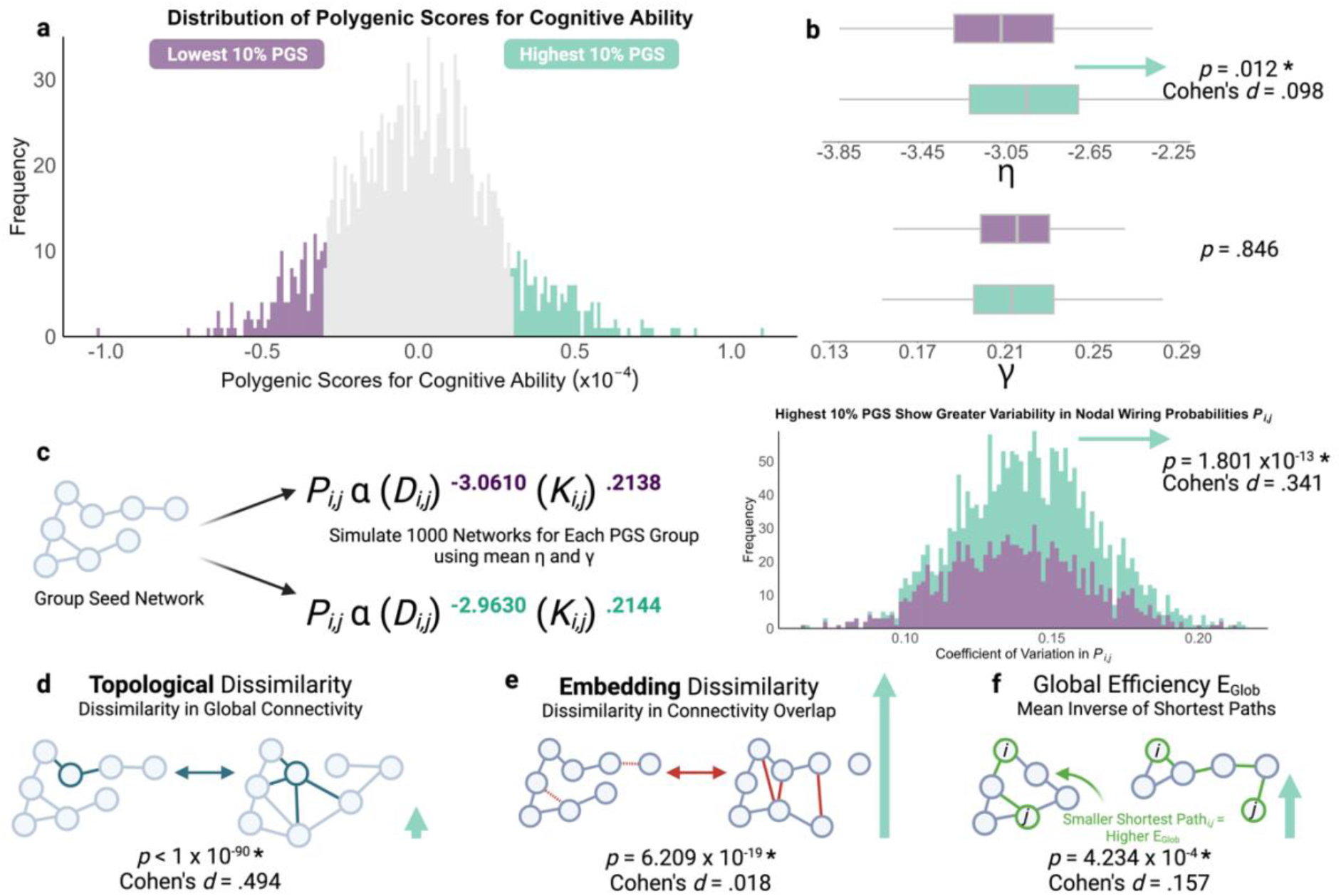
Compared to the bottom 10%, children with the top 10% polygenic scores for cognitive ability display a significantly softer wiring distance parameter η, resulting in more stochastic networks with greater topological dissimilarity, embedding dissimilarity, and global efficiency. **a.** We contrasted children with the lowest (purple) and highest (green) 10% polygenic scores for cognitive ability, resulting in 147 children per group. **b.** Box plot for mean η and γ for each PGS group. **c.** For each PGS group, we simulated 1000 networks using the neighbours generative rule and group mean η-γ parameters. Children with the largest genetic propensity for cognitive ability displayed a significantly greater frequency of more variable nodal wiring probabilities, with variability quantified as the coefficient of variability. This produces networks significantly higher in 3 measures. **d.** The first is topological dissimilarity, describing differences in distributions of local nodal properties. For example, the node in green has differing topology and length of connections between the two networks, resulting in different topology. **e.** The second is embedding dissimilarity, describing differences in the spatial distribution of edges between networks. Differing edges between networks are highlighted in red. **f.** The final measure is global efficiency, a proxy of capacity for information transfer in a network. Smaller shortest paths between any two nodes contribute to higher global efficiency, and larger shortest paths to lower global efficiency. Vertical green arrows for **d-f** are proportional to effect sizes.

How does this softer distance penalty shape the generative process? Logically, networks that form with a weaker η will show significantly more *stochasticity,* the moment-to-moment variability with which connections form. This is because there is slightly less constraint on where longer-range connections can form. In turn, this should produce more variable outcomes, with connections forming in different places across simulations. We tested this with a simple experiment. As shown in **Figure 4c**, taking the average η and γ parameters for each extreme group, and our original seed network, we simulated 1000 networks per parameter combination with the probabilistic GNM. This provided a direct way of testing for the impact of this difference in wiring parameters on the generative process. The softening of the distance penalty, as seen in those with the highest genetic propensity, does indeed increase stochasticity. The variability in the wiring probabilities across the generative process is significantly higher with the softer η parameter (*p* = 1.801 x 10^-13^, Cohen’s *d* = 0.341).

There are two ways of considering the diversity of the resulting networks. The first is topological dissimilarity (**Figure 4d**): are node degree values similarly distributed across simulations? The second is embedding dissimilarity (**Figure 4e**): are connections in the same locations across simulations? The softer distance penalty significantly increases the embedding dissimilarity (*p* < 10^-90^, Cohen’s *d* = 0.494), but has an extremely small effect of reducing topological dissimilarity (*p* = 6.209 x 10^-19^, Cohen’s *d* = 0.018). In other words, when the distance penalty is softer this subtly increases the stochasticity of the generative process, resulting in more structurally diverse outcomes. Finally, does this more stochastic generative process produce more efficient networks? Yes it does: as shown in **Figure 4f**, global efficiency is subtly but significantly higher when the distance penalty is softer (*p* = 4.234 x 10^-4^, Cohen’s *d* = 0.157).

### GNM Parameters and PGSs for Cognitive Ability Significantly Explain Linear Variation in Cognition

Do genetic and generative mechanisms relate directly to cognition across the full sample? To answer this, we tested which linear combinations of individual PGSs, η, and γ parameters accounted for the most variance in cognitive ability (**Figure 5a**), controlling for sex, age, scanner site, and in-scanner motion. We extracted 3 significant components onto which all 3 predictors significantly loaded, evidenced by non-zero CIs, and together accounted for 4.164% of variance (adjusted R^2^) in cognitive ability. η (Mean_XLoading_ = 10.472, 95% CI [10.381 10.563]), γ (Mean_XLoading_ = 13.546, 95% CI [13.471, 13.658)]), and cognitive ability PGSs (Mean_XLoading_ = 34.844, 95% CI [34.806, 34.883]) all loaded positively onto the first component accounting for 4.121% of variance in cognitive ability (Mean_YLoading_ = 7.873, 95% CI [7.854, 7.893], *p_perm_* < .001). Therefore, the first component represents shared variance between computational and genomics predictors of cognition. The second component accounted for .042% of variance in cognitive ability (Mean_YLoading_ = .803, 95% CI [.798 .807], *p_perm_* < .001), where η (Mean_XLoading_ = −31.264, 95% CI [-31.369, −31.158]) and γ (Mean_XLoading_ = −18.470, 95% CI [-18.634, - 18.307]) loaded negatively, and cognitive ability PGS loaded positively (Mean_XLoading_ = 11.562, 95% CI [11.454, 11.669]). Therefore, the second component represents variance in cognitive ability captured by PGSs, relative to η and γ. The third component accounted for the remaining variance (< .001) in cognitive ability (Mean_YLoading_ = .080, 95% CI [.079, .081], *p_perm_* = .006), where γ loaded positively (Mean_XLoading_ = 28.550, 95% CI [28.448, 28.653]), whilst η (Mean_XLoading_ = −15.578, 95% CI [-15.753, - 15.403]) and cognitive ability PGSs (Mean_XLoading_ = −6.053, 95% CI [-6.180, −5.927]) loaded negatively. Therefore, the third component represents remaining variance in cognitive ability captured best by γ. See **Supplementary Table 8** for model performance across all 3 PLSs.

**Fig. 5.**
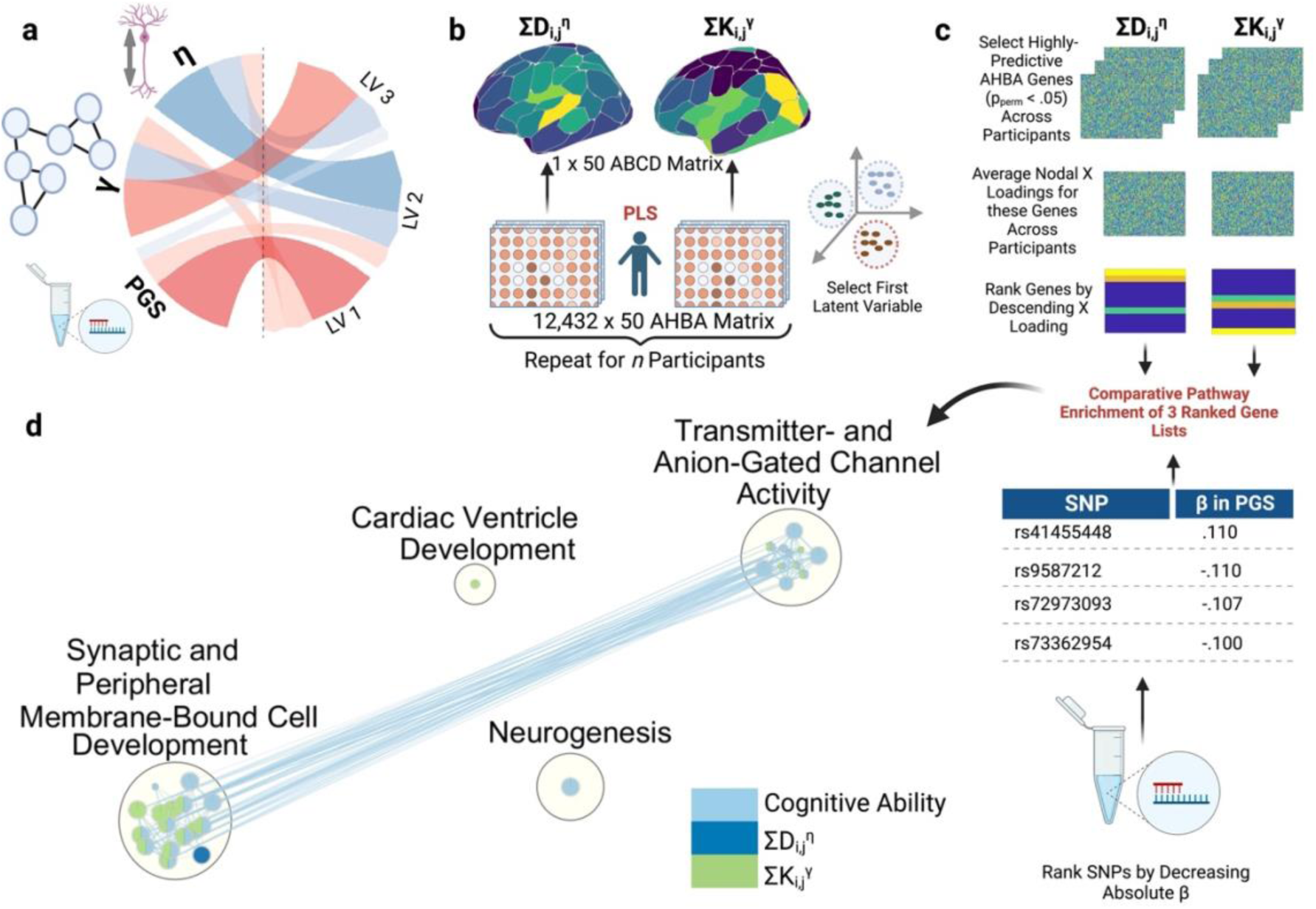
Triangulating the genomic basis of parameterised nodal η, γ, and cognitive ability PGSs. **a** Latent variable (LV) loadings of η, γ, and cognitive ability polygenic scores (PGS), derived from a partial least squares (PLS) regression across 1461 Adolescent Brain Cognitive Development (ABCD) participants. Positive loadings are shown in red, and negative in blue. The width of each arrow represents loading magnitude. **b** For each participant, two PLSs are conducted, using RNA-sequencing genomic data from the left hemisphere of the Allen Human Brain Atlas (AHBA) as predictors of parameterised nodal wiring costs or values, respectively, constrained to the left hemisphere. 10,000 permutations generate a null distribution for each gene’s ability to predict parameterised nodal wiring costs and values, respectively. **c** AHBA genes with a permuted *p*-value less than .05 across all participants are extracted, their nodal X-loadings on the first LV averaged across participants and ranked by decreasing loading. Short nucleotide polymorphisms (SNPs) from the cognitive ability polygenic scores are ranked by decreasing contributions (absolute β). This generates 3 ranked gene lists. **d** To compare the genomic bases of the 3 lists, comparative gene ontology analysis is performed using g:Profiler, significant at *p* < .01. Parameterised nodal wiring value and cognitive ability converge on gene ontologies describing synaptic, peripheral, and membrane-bound cell development. Parameterised nodal wiring costs also share the same cluster, but not nodes, and therefore make an independent contribution. Cognitive ability PGSs only converge with parameterised nodal wiring value, the greatest converge of which is in cell development. The size of each node represents the number of genes in that ontology.

### Overlapping Gene Ontologies for PGS and Structural Brain Organisation Parameters

As linear combinations of η, γ, and PGSs significantly predicted linear variation in cognition, this raises the question of whether the three predictors share a cellular or molecular basis. If they do, then we would expect the genomic correlates of η, γ, and PGSs to converge on common biological aetiologies. To investigate this, we first examined genomic predictors of parameterised nodal wiring costs. For each participant, we conducted two PLSs using RNA-sequencing regional gene expression matrices from the Allen Human Brain Atlas (AHBA), with parameterised nodal wiring costs or value, respectively, as the response variable (**Figure 5b**). To maximise spatial coverage, we constrained our analyses to the left hemisphere, across which all 6 AHBA donors had data. For each AHBA gene, PLS, and participant, we conducted 10,000 permutations to generate a permuted *p*-value. We extracted AHBA genes which significantly (*p_perm_* < .05) predicted either parameterised nodal wiring costs or value across all participants, averaged their nodal loadings onto the first PLS latent variable across participants, and ordered the genes by descending loading (**Figure 5c**). This produced ranked lists of 951 AHBA genes for parameterised nodal wiring cost, and 561 for parameterised nodal wiring value (Akarca et al., 2021; Whitaker et al., 2016). To examine the functional roles of these genes, we performed ordered pathway enrichment analysis for each list separately, using g:Profiler (Kolberg et al., 2020; Raudvere et al., 2019; Reimand et al., 2019). Pathway enrichment analysis tests whether inputted genes are significantly more likely to be grouped together compared to chance. In the case of ordered gene lists, the algorithm searches for the largest sub-list of genes significantly associated with an ontology, and adjusts for multiple comparisons of interdependent gene ontologies (Raudvere et al., 2019; Reimand et al., 2019).

Parameterised nodal wiring costs were significantly enriched for 19 pathways, most of which linked to molecular functions (73.68%). The strongest enrichments were for genes associated with transmembrane transporter activity of metal ions (*p_adj_* = 2.116 x 10^-4^), cations (*p_adj_* = 8.192 x 10^-4^), and inorganic molecules (*p_adj_* = .004). Parameterised nodal wiring value were significantly enriched for 41 pathways, most of which linked to biological processes (43.90%). Rather than transporter activity, the strongest enrichments were for genes associated with different modes of signaling, including synaptic (*p_adj_* = 1.130 x 10^-5^), cell-cell (*p_adj_* = 1.949 x 10^-5^), and anterograde trans-synaptic (*p_adj_* = 2.348 x 10^-5^).

Next, we submitted all 76,745 SNPs in the PGS, ranked by descending absolute β, to pathway enrichment analysis. SNPs predictive of cognitive ability were significantly enriched for 297 pathways, mostly encoding biological processes (49.50%). The strongest enrichments were for genes associated with synaptic (*p_adj_* = 3.485 x 10^-29^) and post-synaptic (*p_adj_* = 3.684 x 10^-26^) membranes, alongside synaptic (*p_adj_* = 7.602 x 10^-20^) and trans-synaptic (*p_adj_*= 3.821 x 10^-20^) signaling. Summaries for all pathway enrichment analyses and g:Profiler links are provided in **Supplementary Table 9**.

To assess possible shared ontologies of η, γ, and PGSs, we submitted all three ranked gene lists to a multi-query comparative pathway enrichment analysis (https://biit.cs.ut.ee/gplink/l/h44jDUAPRv, **Supplementary Table 10**). This produced 3 adjusted p-values for each pathway, which represents the likelihood that each gene list was significantly enriched compared to chance. 96 gene ontologies were significantly enriched for at least one predictor (*p_adj_*< .01), and mostly for molecular functions (43.75%).

When grouped on connectivity strength, 4 ontological groups emerged (**Figure 5d**), 2 of which were shared by all predictors. Greater convergence between parameterised nodal wiring value and cognitive ability PGSs occurred for synaptic, peripheral, and membrane-bound cell development, and less for transmitter-and anion-gated channel activity. Whilst parameterised nodal wiring cost also contributed to synaptic, peripheral, and membrane-bound cell development; it made a unique contribution towards presynaptic development. Cognitive ability PGSs were distinguished by neurogenesis, and parameterised nodal wiring value by cardiac ventricle development.

## DISCUSSION

Human brain development proceeds via dynamic interactions between our unique genetic background, our experience, and physical constraints on the formation of complex neuronal networks (Gottlieb, 2007; Kaiser & Hilgetag, 2004; Westermann et al., 2007). One popular way of conceptualising those physical constraints is as an economic trade-off between the metabolic cost of creating or maintaining new connections, and the topological value those connections provide (Akarca et al., 2021, 2022; Betzel et al., 2016; Bullmore & Sporns, 2012; Carozza et al., 2022; Vértes et al., 2012; Zhang et al., 2021). In this framework, structural brain development can be conceptualised as a negotiated trade-off, maximising topological value whilst minimising cost. We investigated whether and how population-level variability in this trade-off is related to known genetic variants associated with cognitive performance (Savage et al., 2018). Across 2154 children, a homophily-based model provided the best account of structural brain organisation, indicating that across development, topologically valuable connections are those that form between nodes with similar connectivity profiles. This trade-off differs according to someone’s polygenic propensity for cognitive ability. At the extremes this is most apparent as a softening of the distance penalty, giving rise to more stochastic, diverse and efficient networks for those with the highest PGS. Importantly, across the whole sample, the two parameters that govern the trade-off at an individual level (η, γ) share variance with known genetic variants as they predict cognitive ability. A comparative pathway enrichment analysis shows that these known variants converged on common genetic ontologies with wiring costs η and value γ. Put simply, population-level genetic variability that is linked to cognitive performance overlaps with variability in generative wiring parameters. One possible reason is because of shared cellular and molecular mechanisms, with regional co-expression with topologically valuable areas.

In line with prior work across species, scales and neurodevelopmental conditions (Akarca et al., 2021, 2022; Betzel et al., 2016; Carozza et al., 2022; Zhang et al., 2021), we found a generative model combining wiring cost and homophilic value most accurately recapitulated structural brain development in children. This suggests that prioritising forming edges between nodes with shared neighbours is a major organisational principle for childhood structural brain development. This reflects a key organisational principle for economical brain development, whereby topological neighbours are also likely to be anatomical neighbours (see Bullmore & Sporns, 2012). Interestingly, our finding also aligns with network neuroscience accounts of general cognitive ability (Barbey, 2018) emerging from a trade-off between minimising wiring cost, which prefers local efficiency, with maximising wiring topology, which prefers global efficiency.

Having a high genetic propensity for cognitive ability was significantly associated with a weaker distance penalty, resulting in more stochastic, diverse and globally efficient networks. Stochasticity in neural networks, particularly in terms of wiring, emerges across multiple scales, ranging from genetic splicing, epigenetic chromatin modifications, and differing numbers of cell adhesion and intra-cellular signalling molecules steering neuronal growth cones to build a connectome (Honegger & de Bivort, 2018). This stochasticity may be evolutionarily adaptive in several ways (Honegger & de Bivort, 2018), such that sufficient variation in the population allows members to display adaptive behaviours which are either stable (diversified bet-hedging) or vary (phenotypic plasticity), and this requires fewer successive generations to achieve (gene saving) compared to reduced stochasticity. We show that common genomic variation in the population associated with cognitive ability encodes greater stochasticity, resulting in more efficient networks, which may offer adaptive advantages. Likewise, recent work (Carozza et al., 2023) has shown that the generative trade-off can be shifted when organisms develop in unpredictable circumstances, again introducing stochasticity into network formation that may have adaptive advantages. Our findings show that relatively small, albeit significant, changes in the wiring parameters can have a disproportionate impact on network stochasticity and diversity, presumably because the process of network formation of probabilistic, so even small shifts in economic constraints will cascade across the generative process.

The specificity of the relationship between genes, brain development and cognitive ability depends on variability in genetic relatedness and the influence of individual genes. At the extreme of genetic relatedness, twin studies have demonstrated genetic influences on specific aspects of network topology, such as rich-club nodes with weak peripheral edges, as well as inter-hub connectivity (Arnatkeviciute et al., 2021). At the extreme of genetic influence, high-impact single-gene mutations, implicated in neurogenetic disorders, exhibit tight spatial coupling with altered structural brain organisation. Transcriptional vulnerability suggests that differences in regional gene expression profiles shape ongoing structural brain development, with knock-on implications for cognition. A prior study validated this model in participants with neurogenetic conditions, whereby patterns of morphometric dissimilarity were most strongly associated with copy number variation, and the spatial patterns could be recovered in gene expression maps derived from control post-mortems (Seidlitz et al., 2020). Our findings reflect the opposite end of the spectrum – common genetic variation in non-related neurotypical children. Thus, inevitably the overall strength of the genetic influences are smaller because they reflect population-level variation, rather than that between twins or following a high-penetrance mutation.

We examined the link between wiring stochasticity and genetic propensity for cognitive ability using two approaches. The first demonstrated greater stochasticity through a weaker distance penalty in children with a higher genetic propensity for cognitive ability, using children at the extremes of the PGS distribution. The second used a comparative gene enrichment for wiring parameters and PGSs across the entire sample to establish the points of intersection between these predictors. Across the entire sample, enriched cellular and biological processes overlap with those associated with cognition-linked common variants, and the overlap is greater with the ‘value’ part of the equation. This suggests that genetic variations most predictive of cognitive ability align more closely with genes most predictive of wiring *updating*, rather than fixed cost. One reason for this may be greater variability in nodal wiring value and topology compared to wiring distance. From a biological standpoint, wiring cost is determined by distance, with set metabolic costs, and is physically constrained. Therefore, stochasticity in a GNM framework, or variability in the likelihood of two nodes being connected, at a population-level, reflects variability in wiring value more so than cost. This overarching association of stochasticity with wiring value occurs *in tandem* with associations between stochasticity and wiring cost in a subset of participants at the extreme tails of a genotypic, namely PGS, distribution.

Whilst the current study was the first to triangulate cognition, participant-derived and donor-derived genetics with structural brain development metrics at a population level, there are inevitable limitations. First, our sample is slightly overrepresented for children with parents with higher educational backgrounds and of European and Asian descent. Second, we included participants with high-quality neuroimaging data which could skew our sample towards those with low levels of hyperactivity and associated head motion. Third, we established genomic patterning of GNM parameters using adult post-mortem atlases. Whilst recent efforts have been made to collate post-mortem gene expression data across the lifespan (Jiao et al., 2019), including childhood, these offer considerably less spatial resolution than the AHBA (see work by Arnatkevičiūtė and colleagues (2019) for further discussion of AHBA limitations). Third, previous work has suggested alternative generative modelling specifications, which may challenge the conclusion that homophily based models perform best. For example, a variation accounting for change in topology as part of a growth model outperformed static models (Oldham et al., 2022), such as those used in the current study. Further, explicitly incorporating genetic constraints improved model performance, which otherwise fails to recapitulate spatial topography in absence of a seed network (Arnatkeviciute et al., 2021; Oldham et al., 2022). Finally, the current GNM specification is binary, with a single pair of parameters describing global development, adding a single edge at each iteration, and assumes that the morphological and microstructural properties of these connections are consistent across development and the cortex. Inevitably, synaptic connections are not added in a binary fashion but are gradually strengthened or weakened through experience-dependent synaptic plasticity, and in other cases completely pruned (Huttenlocher & Dabholkar, 1997). Advances in the complexity of generative models will be needed to capture this nuance.

In summary, trade-offs between nodal wiring cost and value accurately recapitulated the structural brain development properties of 2153 children. Focusing on a subset of children at the extreme tails of a distribution for polygenic scores revealed a weaker distance penalty, which has a disproportionate impact on network stochasticity, diversity and efficiency. Across the entire sample, common genomic variation associated with cognitive ability overlapped most with wiring value constraints. Together, we demonstrate how shared underlying cellular and biological processes begin to shape variability in structural brain development and cognitive ability in childhood.

## METHODS

### Participants

The scale and depth of the Adolescent Brain Cognitive Development (ABCD) Study provides an unparalleled opportunity to integrate computational models of structural brain development, genomic variation, and population-level variation on cognitive ability. This prospective longitudinal dataset charts multi-modal brain and cognitive development of 11,878 genotyped US children, with baseline data collected at 9-11 years old. Beyond considerable statistical power, the ABCD sample benefits from high external validity due to heterogeneity in ethnicity, socio-economic status, and sex, with 21 recruitment centres distributed across the US (Casey et al., 2018; Karcher & Barch, 2021).

### Cognition

To measure the structure and variance of cognitive scores, we selected participants corresponding to the baseline assessment from the ABCD Study Data Release 4.0 (DOI: 10.15154/1523041), comprised of 11876 children (*N* = 5668 male), aged between 8.92 and 11.08 years (Mean = 9.92 ± .63 years), recruited from 21 sites across the United States. Overall cognition, measured by the National Institutes for Health Toolbox Total Composite Cognitive Score (NIH-TB-Comp), varied considerably, between 32 and 221 (Mean = 100.40 ± 18.02). The ABCD Study was designed to reflect the US population composition (Karcher & Barch, 2021) through race (52.4% White, 17.3% Hispanic, 15.1% Black, 2.1% Asian, and 13.2% Other), and socio-economic status (SES) distribution, including parental working status (69.6% Working, 4.3% Looking for Work, .6% Retired, 17.4% Stay-At-Home Parents, and 8.1% Other), highest parental educational attainment (6.6% Less than 12^th^ Grade, 10.6% High School Graduates, 16.5% with some College Experience, 13.0% with an Associate’s Degree, 28.1% with a Bachelor’s Degree, 22.0% with a Master’s Degree, and 3.2% with a Doctoral Degree), and income-to-needs ratio (INR) (M= 3.36 ± 2.61). Ethical approval for most recruitment sites was obtained from the central Institutional Review Board (IRB) at the University of California, with the remaining sites granted local IRB approval (Auchter et al., 2018).

### Neuroimaging

From the participants described above, we selected a stratified subset of 2193 genotyped children who had high-quality T1-weighted (T1w) MRI, rsfMRI, and single-run DWI data. To ensure that our subsample was representative of the larger ABCD cohort, we stratified based on 3 variables: general cognitive ability, measured by the National Institutes of Health Toolbox for the Assessment of Neurological and Behavioural Function (NIH-TB) Composite Total Cognitive Age-Corrected Score, and binned in four groups (32-88, 88-100, 100-112, 112-221); age at time of MRI scans (months), binned into two groups (107-119, 119-133 months); and parent-reported sex (2 levels – male and female). These and other demographic variables are visualised in **Supplementary Figure 3**.

From the stratified sample described above, we successfully processed data from 2154 participants across 3 parcellations: Schaefer 100-node 17-network (Schaefer et al., 2018), Brainnetome 246-node (Fan et al., 2016), and Schaefer 400-node 17-network parcellations (Schaefer et al., 2018). A series of bootstrapped Kolmogorov-Smirnov tests demonstrated no significant difference between the distributions of the successfully processed participants and the larger ABCD sample in terms of age [*D*(2154) = .016, *p* = .572], NIH-TB-Comp age-corrected scores [*D*(2154) = .012, *p* = .955], or INR [*D*(2154) = .028, *p* = .090]. Further, a 2×2 and 7×2 Chi-Square test revealed no significant differences in distributions of sex [X^2^ (1, 2154) = .027, *p* = .869] and parental working status [X^2^ (4, 2154) = 2.867, *p* = .580], respectively. However, a 7 x 2 Chi-Square test and 5 x 2 Chi-Square test revealed significant differences in distributions of highest parental education [X^2^ (6, 2154) = 19.767, *p* = .003, *OR* = 1.374] and race [X^2^ (4, 2154) = 16.377, *p* = .003, *OR* = 1.335], respectively. Note that whilst we initially included participants who had successfully processed data for the Gordon 333-node parcellation (Gordon et al., 2016), we later removed this requirement as our downstream connectome thresholding procedure (Betzel et al., 2019) required an equal number of nodes.

To calculate cognitive ability PGSs, we selected genotyped participants from the ABCD Baseline with complete demographic data and excluded those with axiom plate 461 known to be problematic, generating N = 10,979. Note that we downloaded genetic data from the ABCD Study Data Release 3.0, which was automatically provided by the most recent data release, due to no differences between releases. To use cognitive ability PGSs as a predictor for GNM wiring parameters, we retained participants who’d passed genomic QC and had successfully processed structural connectomes (N = 1,468).

### Neuroimaging Acquisition and Processing

We generated SC matrices for each participant across 3 parcellations of varying spatial resolution, as described above, for which we conducted group-level analyses. Note that the following sections contain information from the QSIprep boilerplate.

### Structural MRI

We downloaded high-quality Fast-Track structural MRI and DWI from the ABCD-BIDS Community Collection (https://collection3165.readthedocs.io/en/stable/). T1w images were acquired during a single session across an axial plane, with a multiband gradient echo sequence, 8° flip angle, 256 x 256 matrix size, 1060ms inversion time, and 1mm isotropic resolution (Casey et al., 2018; Hagler et al., 2019). Since the ABCD study is a collaborative effort spanning 21 acquisition sites, participants were scanned using of three possible 3-Tesla scanners, using the following additional scanner-specific parameters: Siemens Prisma VE11B-C (Repetition Time (TR) = 2500ms, Echo Time (TE) = 2.88ms), with 176 slices, General Electric (GE) MR750 DV25-26 (TR = 2500ms, TE = 2ms) with 208 slices, and Philips Achieva dStream or Ingenia (TR = 6.31ms, TE = 2.9ms) with 225 slices. For further details, see Casey et al., 2018 and Hagler et al., 2019.

We processed structural MRI data using QSIprep 0.15.3 (Cieslak et al., 2021), implemented through *Nipype 1.7.0* (Gorgolewski et al., 2011), *Nilearn 0.9.0* (Abraham et al., 2014) and *Dipy 1.4.1* (Garyfallidis et al., 2014). Following the T1w image corrected for intensity non-uniformity using the N4 Bias Field Correction algorithm (Tustison et al., 2010), the image was skull-stripped using The Open Access Series of Imaging Studies (OASIS) as a target, creating a T1w reference image. Volume from the T1w reference and OASIS target were used to non-linear spatially register and normalise the T1w reference to the ICBM 152 Nonlinear Asymmetrical template version 2009c, as a normative brain template for children aged between 4.5 and 18.5 years old (Fonov et al., 2011). The normalised image was then segmented into cerebrospinal fluid, cortical gray matter and WM using FMRIB’s Automated Segmentation Tool (ANTs; Zhang et al., 2001).

### DWI

Across all 3 scanners, during a single session 81 DWI slices of size 140 x 140 voxels were acquired with 4 b-shells (500, 1000, 2000, 3000 s/mm^2^) across a total of 96 diffusion directions (b = 500 s/mm^2^, 6 directions; b = 1000 s/mm^2^, 15 directions; b = 2000 s/mm^2^, 15 directions; b = 3000 s/mm^2^, 60 directions), with 1.7 mm isotropic resolution, and MultiBand Acceleration Factor 3 (Casey et al., 2018; Hagler et al., 2019). In addition, the following scanner-specific parameters were used: Siemens (TR = 4100ms, TE = 88ms, 90° flip angle), GE (TR = 4100ms, TE = 81.9ms, 77° flip angle), and Philips (TR = 5300ms, TE = 89ms, 78° flip angle).

Using QSIprep 0.15.3, DWI data underwent resampling to T1w space, intensity normalisation through being scaled by b=0 means, linear co-registration to T1w using ANTs, denoising, and susceptibility distortion (Eddy currents) correction using FSL TOPUP (Andersson et al., 2003; Smith et al., 2004). The processed DWI data was reconstructed using DSI Studio, which uses non-parametric model-based approaches to fit a multi-tensor model, with generalized q-sampling imaging orientation distribution functions (ODFs) (Yeh et al., 2010) estimated with mean diffusion distance ratio of 1.25. The DWI data then underwent deterministic tractography (Yeh et al., 2013) with 5 million randomly-seeded streamlines, between 30mm and 250mm in length, and 1mm step size. We chose deterministic tractography based on evidence of more accurate connectome reconstructions and lower false-positive rates compared to probabilistic tractography (Sarwar et al., 2019). Note, however, that deterministic tractography does not provide an estimate of confidence of tract orientation, unlike probabilistic tractography.

### Connectome Construction

We considered structural connectivity as the number of streamlines terminating at each node, yielding a *nroi* x *nroi* matrix per participant, where *nroi* denotes the number of regions of interest in each of the 3 parcellations investigated. We then averaged across SC for all participants for each parcellation, yielding a single *nroi* x *nroi* matrix. We conducted two sets of thresholding procedures. First, to remove spurious connections, we applied a 60% consensus threshold to all individual connectomes. Second, using these connectomes, we applied a consensus-based distance-dependent threshold across all participants, which has been shown to preserve key statistical network properties and edge length distributions (Betzel et al., 2019), yielding a single consensus network for all participants, acting as the group target connectome. This connectome in the Schaefer 100-node parcellation was plotted in **Figure 1b** using the brainconn R package (Orchard et al., 2021). Third, to construct the seed for the group models, we selected connections present in at least 95% of participants. After running the group models, we conducted additional thresholding at the individual level, namely streamline thresholding at 27 streamlines to produce individual target connectomes, with the seed the same as the group model. At the group-level, 608.08 new edges were added to the seed to reach the target. At the individual-level, an average of 594 (± 49.42) connections were added.

### ABCD Genomics Acquisition and Quality Control

Genotyping data was provided by the ABCD study, where short nucleotide polymorphisms (SNPs) from saliva and whole blood samples were extracted and processed by Rutgers RUCDR, and assayed using the Affymetrix NIDA SmokeScreen Array, made up of 646,247 markers (Baurley et al., 2016; Uban et al., 2018). Genotyping quality control was conducted by the ABCD DAIRC Team, using recommendations from the Ricopili pipeline which, in brief, consisted of pre-imputation quality control to ensure consistency of sample identifiers, PCA to identify duplicates and population structure, imputation to a reference panel to maximise SNPs, alongside post-imputation clumping of genome-wide significant SNPs and incorporation of covariates (see Lam et al., 2020; Uban et al., 2018). Together, 11099 participants with 516,598 SNPs passed ABCD QC (see ABCD Study 4.0 Release Notes).

Our discovery GWAS dataset was provided by Savage and colleagues (2018), which consisted of 267,867, principally European, participants, using the NCBI GRCh37 Genome Build, spanning 9,295,118 SNPs. As part of QC for the discovery dataset, we removed SNPs with minor allele frequency (MAF) less than .10, imputation information scores less than .80, ambiguous and duplicated SNPs with multiple or missing bases, resulting in 6,101,992 SNPs.

Our target dataset consisted of 10,979 ABCD participants with high-quality genotypes and complete demographic information. Using PLINK 1.9 (Chang et al., 2015; Purcell et al., 2007; Purcell & Chang, 2019), we performed additional QC on the target dataset, detailed elsewhere (Choi et al., 2020). 8320 SNPs had incorrectly formatted unique identifiers, therefore associated allelic data was obtained from the Ensembl Gene Browser (GRCh37), and non-ambiguous, unique matches used to update the binary PLINK files. Next, we removed SNPs with MAF < .10, Hardy-Weinberg Equilibrium < 1e^-6^, genotyping rate > .99, and missingness rate < .01, and then pruned the resulting SNPs with linkage disequilibrium variance (LD r^2^) higher than .25 in steps of 50 variants across a window size of 200 variants. Further, we removed participants with heterozygosity F-coefficients greater than 3 standard deviations above or below the mean. We retained effect alleles which did not require strand flipping, recoding, or whose bases were not complementary between the base and target datasets. Finally, we removed participants with discordant sex information and relatedness > .125. Together, 6528 participants with 197,798 SNPs passed QC.

### Polygenic Scores

PGSs summarise the number of SNPs contributing towards a phenotype, each weighted by effect sizes from a discovery dataset. Since ancestry is a major confounding variable in PSs (see Choi et al., 2020), we first inferred the underlying population structure using k = 4 ancestral groups in fastStructure (Raj et al., 2014). We found that the optimal k ranged between 1 and 4, and therefore, for simplicity, decided upon k = 2 clusters. Following prior ABCD studies (Loughnan et al., 2021), we stratified our analyses into those of over 90% European ancestry (N = 7500) and classified the remaining participants (N = 3479) as Non-European. We performed the PS in PLINK 1.9. We clumped all SNPs and retained those with a LD r^2^ > .1 and calculated the PS against 7 p-value thresholds for SNP inclusion (.001, .05, .1, .2, .3, .4, .5). Finally, to account for population structure, we adjusted our PS by including 6 population principal components as covariates. This resulted in PGSs calculated for 6528 participants, 1486 of which overlapped with the stratified sample, and 1461 of which had successfully processed structural connectomes and GNMs.

### Allen Human Brain Atlas Microarray Processing and Parcellation Choice

We used the Allen Human Brain Atlas microarray data as measures of regional brain gene expression (https://human.brain-map.org/). This dataset contains transcriptomic data for 6 donors (*N* = 5 male) aged between 24 and 57 years old, across 3 ethnicities (AHBA Case Qualification and Donor Profile Technical White Paper), spanning over 62,000 probes and 26,000 genes, with approximately 500 samples collected per hemisphere per donor (AHBA Microarray Survey Technical White Paper). To reduce the influence of methodological factors, AHBA performed additional within-batch quality control, such as adjusting for intensity distribution, RNA quality, batch effects, and dissection method, and between-batch quality control steps, including aligning human brain atlas control and mean expression values (AHBA Microarray Data Normalization Technical White Paper). Further methodological details are available from the AHBA Technical White Papers listed above (https://help.brain-map.org/display/humanbrain/Documentation).

Whilst we initially intended to use the Schaefer 400-node parcellation for genetic and neuroimaging analyses, to maximise spatial resolution, we found that the distribution of the number of samples provided for each AHBA region [Mean = 8.94, Median = 6.50, Mode = 1.00] and the number of donors providing samples [Mean = 3.39, Median = 2.00, Mode = 2.00] was positively skewed, suggesting that a lower-resolution parcellation may offer better spatial coverage (**Supplementary Figure 1**). Further, the degree of bilateral missingness decreased from 13.00% for the Schaefer 400-node parcellation, to 3.66% for the Brainnetome 246-node parcellation, to 0% for the Schaefer 100-node parcellation, with RNA-sequencing probe selection, 0.5 intensity-based filtering, and 2mm region assignment distance threshold. Therefore, we selected the Schaefer 100-node parcellation for individual-level analyses.

Although within-batch and between-batch normalization was conducted by the AHBA, additional variability can occur through researcher-led decisions about assigning AHBA gene expression values to a parcellation. Therefore, we followed the best-practice guidance of Arnatkevičiūtė and colleagues (2019) when parcellating the AHBA microarray data, and used the *abagen* toolbox (Arnatkevičiūtė et al., 2019; Hawrylycz et al., 2012; Markello et al., 2021) in Python 3.10. In brief, probes were reannotated with up-to-date genetic labels from Arnatkevičiūtė and colleagues (2019), and intensity-based filtering performed, such that only probes whose expression exceeded background noise in at least 50% of brain regions were retained. In instances where multiple probes mapped onto the same gene, the probe with the highest correlation in RNA-sequencing data between the two donors with such data were selected. Note that whilst different probe selection approaches exist, RNA-sequencing arguably offers the highest validity, due to reduced noise and lack of reliance on known genetic associations (Arnatkevičiūtė et al., 2019). However, since only two donors have such data, we decided to use RNA-sequencing for probe selection and as an external reference for the microarray data, for which all 6 donors have data. The MNI coordinates of the AHBA samples were then updated using the *alleninf* Python package (Gorgolewski et al., 2014), and mapped to each voxel in the Schaefer 100-node parcellation with a distance threshold of 2mm. Regional gene expression values for each donor were then normalized using a scaled robust sigmoid function previously shown to be robust to outliers (Arnatkevičiūtė et al., 2019; Fulcher & Fornito, 2016). To ensure consistency of gene expression values with RNA-sequencing, we excluded microarray genes absent in the RNA-sequencing matrices and whose Spearman’s correlation with RNA-sequencing gene expression exceeded .75. We then averaged across donors to produce a 100 (number of nodes) x 12431 (number of genes) matrix. We restricted our analyses to the left hemisphere, for which all 6 donors had data, yielding a dense 50 x 12431 matrix.

### Cognitive Measures

The NIH-TB was administered to all participants via touchscreen, used to measure several broad cognitive domains spanning attention, working memory, language, and executive functioning, previously validated across different racial groups, educational levels, and ages (Weintraub et al., 2013). A total cognition composite score, derived from averaged standardised raw scores from the 7 subscales, provided a measure of combined crystallised and fluid intelligence, and hence overall cognition (Akshoomoff et al., 2013). We used age-corrected scores for our analyses. Cognitive subscales are detailed in **Supplementary Table 1**.

### SES

We quantified SES as INR, parental highest educational attainment, and parental working status. We calculated income-to-needs ratio by dividing the median of each total household income bin by the 2018 US Federal Poverty Line for the corresponding household size. A smaller ratio indicated greater degree of poverty, with a ratio of 1 signifying the poverty line. Parental highest educational attainment and working status were initially coded by ABCD as 21 and 13 discrete levels, respectively. To simplify our stratification, we recoded these two variables into 7 and 5 new levels, respectively. SES measurement scales are further described by Barch and colleagues (2021).

### Generative Network Models

We used the Brain Connectivity Toolbox (Rubinov & Sporns, 2010) in MATLAB R2022a (The MathWorks, Inc., 2022) to construct GNMs. GNMs simulate connectivity from a sparse seed network by iteratively calculating the probability of any two nodes being wired (**Equation 1**), until the observed (empirical) connectome is reached (Betzel et al., 2016; Kaiser & Hilgetag, 2004; Vértes et al., 2012).

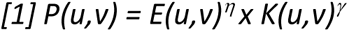

The GNM is made up of a geometric and a non-geometric term. The geometric term refers to the Euclidean distance between two nodes [*E(u,v)*], scaled by the wiring cost η, and therefore favours short-range metabolically ‘cheap’ connections. Rather than metabolic cost, the non-geometric term suggests that topology drives connection formation, whereby the probability of connection formation between two nodes is given by a topological term [*K(u,v)*] scaled by the wiring value γ. K represents different network properties driving connectome formation, and is grouped into 4 classes: clustering, degree, homophily, and spatial. The current study considered 5 generative models, corresponding to the best-performing model in each class (average clustering, average degree, geometric), and both homophily models (matching and neighbours), based on prior studies (Akarca et al., 2021; see Betzel et al., 2016 for model specifications).

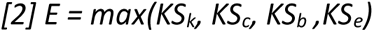

The most common way to evaluate generative model performance is through an energy function (**Equation 2**; Akarca et al., 2021, 2021; Arnatkeviciute et al., 2021; Betzel et al., 2016; Zhang et al., 2021). This is a global measure of model fit. A Kolmogorov-Smirnov (KS) test assesses whether the distribution of 4 statistics significantly differs between the synthetic and observed connectomes: degree (*k*), clustering (*c*), betweenness-centrality (*b*), and edge length (*e*). The largest KS-statistic is the model energy (Betzel et al., 2016). Therefore, the model class with the smallest energy, hence smallest dissimilarity, was deemed the best fit.

We performed two sets of analyses – at the group level, and at the participant level. To establish which of the 5 wiring rules described above best accounts for the wiring properties of the group connectome, and using which parameters, we performed a grid search using 100,000 equally spaced random η [-7 ≤ η ≤ 7] and γ [-7 ≤ γ ≤ 7] combinations (99,856 unique combinations). We selected the η and γ boundaries as in a recent GNM paper (Akarca et al., 2021). To select the best-fitting model, we used 3 criteria. The first criterion was energy, where the best-fitting model should have the lowest energy. Whilst energy is a global measure of fit, we also assessed the ability of the synthetic connectomes to capture distributions of local nodal properties through topological similarity. The final criterion was spatial similarity, which compares the of distribution of nodal edges (see Results) between the synthetic and observed connectomes. We ranked each model according to these 3 criteria and selected the model with the smallest cumulative rank as the best fit. For the best-performing model across all participants, we performed an additional 50,000 simulations for each participant, using η and γ parameter boundaries corresponding to the top 10% lowest energy group simulations (49,729 unique combinations). To explore how GNM terms varied across the simulation, for our individual-level analyses, we collected node-wise parameterised D_i,:_ and K_i,:_ terms for each new connection added until the target was reached. Throughout, we used the ggseg R package (Mowinckel & Vidal-Piñeiro, 2020) to plot nodal measures on the cortical surface, and the ggplot2 R package (Wickham, 2016) to visualise correlations of observed and simulated network statistics.

### Statistical Analyses and Data Processing

We processed cognitive, demographic, and genetic data in R version 4.1.2 (R Core Team, 2022), and conducted statistical analyses in MATLAB R2022a (The MathWorks, Inc., 2022).

### Data Transformations

We coded parents who failed to answer questionnaires or were unsure as missing. We imputed missing values across the 7 cognitive subscales (n = 241), NIH-TB-Comp measure (*n* = 397), age (*n* = 75), race (*n* = 194), INR (*n* = 386), sex (*n* = 75), parental education (*n* = 92) and parental working status (*n* = 131) using predictive mean matching through Multivariate Imputation by Chained Equations, implemented using the ‘mice’ package (van Buuren & Groothuis-Oudshoorn, 2011).

### Principal Component Analysis

Rather than restricting our analyses to specific cognitive subscales, we extracted a general ‘*g*’ factor through principal component analysis (PCA) of the 7 NIH-TB subscales using the FactoMineR package (Lê et al., 2008).

### AHBA Partial Least Squares Regression

To explore the relationship between AHBA gene expression and nodal parameterised wiring value and costs, we conducted two partial least-square regressions (PLSs) for each participant. This is a multivariate data dimensionality-reduction technique which models the weighting or loadings of combinations of predictors that accounts for the maximum variance shared by the predictors and the response variables (Geladi & Kowalski, 1986), similar to a latent variable. PLSs are superior to traditional regressions in that they can cope with many highly correlated predictors and response variables. For each participant, the predictor variables were the left-hemisphere regional AHBA gene expression values, with dimensions 50 (regions) x 12431 (genes). The response variable in PLS1 was nodal parameterised wiring costs (1 x 50 matrix), whilst the response variable in PLS2 was nodal parameterised wiring value (1 x 50 matrix). For each PLS and participant, we conducted 5000 permutations to derive a null distribution of the loadings of each gene onto each of 3 PLS components (as in Akarca et al., 2021).

### Linking Genes, Brain, and Cognition Through Linear Models and PLSs

To explore the relationship between genes, brain, and cognition, we conducted 3 PLSs across participants with the following specification, each with 10-fold cross-validation: η and γ as the two predictors, cognitive ability as the response; η and γ as the two predictors, cognitive ability PGS as the response; η, γ, and cognitive ability PGS as the three predictors, cognitive ability as the response. For each PLS, to test component significance, we conducted 10,000 permutations, and applied the Procrustes rotation to correct for sign-flipping (Bastien, 2008). To test loading significance, we conducted 10,000 bootstraps, from which we extracted mean loadings and 95% confidence intervals (CI). If the 95% CI did not cross zero, the predictor loading was considered significant.

### Gene Ontology and Pathway Enrichment Analyses

To examine possible common biological pathways and correlates of parameterised nodal wiring values, costs, and cognitive ability PGSs, we conducted gene ontology (GO) and pathway enrichment analyses for each of the 3 ranked gene lists. The first list was ranked by decreasing X loading onto the first extracted PLS component of parameterised nodal wiring costs regressed onto the AHBA gene expression matrix across the left hemisphere. The second list was ranked according to the equivalent analysis for parameterised nodal wiring value. The third list contained the 76,745 SNPs which were included in the cognitive ability PGS. We ranked each SNP by decreasing absolute contribution β to the PGS. We conducted pathway enrichment analyses for each of the 3 gene lists, alongside multi-query ranked comparative enrichment tests, using g:Profiler through the gProfiler2 v0.2.1 R package (Kolberg et al., 2020), using the Gene Ontology (v e107_eg54_p17_bf42210; Ashburner et al., 2000; The Gene Ontology Consortium, 2021) database. The results were visualised using EnrichmentMap 3.3 (Merico et al., 2010) in Cytoscape v3.9.1 (Shannon et al., 2003), and annotated using the Cytoscape AutoAnnotate plugin (Kucera et al., 2016) through the Community Cluster algorithm from clusterMaker2 (Morris et al., 2011). The Cytoscape session used to visualise these networks is available in the OSF repository accompanying this paper.

## Code Availability

All MATLAB and R scripts are provided at https://github.com/AlicjaMonaghan/abcd_genes_brain_cognition

## Data Availability

We used data from Release 4 of the ABCD study (http://dx.doi.org/10.15154/1523041). Access derived data at https://osf.io/xf8bc/?view_only=7a4a3e30e4624d7ea5156242f8eac3ff

