## Supplementary Materials for "Population-Level Genetic Variation Shapes Generative Brain Mechanisms"

**Table S1. Description of 7 National Institutes of Health Toolbox Cognitive Assessments administered to Adolescent Brain Cognitive Development (ABCD) Participants**

| NIH-TB Subscale | Cognitive Domain | Description |
| --- | --- | --- |
| Flanker Inhibitory Control and Attention Test (Eriksen & Eriksen, 1974; Rueda et al., 2004) | Attention | Children fixate onto a central cross, after which a set of 5 horizontal stimuli appear, either fish (for children younger than 8 years old) or arrows (for children older than 8 years old), all facing either in the same direction (congruent) or in different directions (incongruent). The child must report the direction of the central stimulus (Weintraub et al., 2013). |
| Dimensional Change Card Sort Test (Zelazo, 2006) | Set-Shifting | Children are presented with two target cards which vary across two dimensions, such as shape and colour. They are then presented with two test cards and asked to sort them according to the first dimension (pre-switch phase) and then the second dimension (post-switch phase). |
| List Sorting Working Memory Test (Tulsky et al., 2014) | Working Memory | During Part 1, children are presented with a list of objects belonging to Category 1 and must repeat these objects sorted from smallest to largest. During Part 2, children are presented with a list of objects belonging to Category 2 and must report objects sorted from smallest to largest, first from Category 1, and then from Category 2. |
| Picture Sequence Memory Test (Dikmen et al., 2014) | Episodic Memory | During each of 3 learning trials, children were presented with a sequence of images of actions, each of which narrated. After presentation of each image, the image would be minimised and return to its position within the sequence. During the test phase, the images were presented in a random spatial order, and the child must reorder them according to the order presented during the learning trials. Difficulty was manipulated by sequence length. |
| Oral Reading Recognition Test (Gershon et al., 2013) | Language | To test pronunciation and orthographic recognition, children were asked to repeat a set of words varying frequency, age of acquisition, frequency of syllables and phonemes (Gershon et al., 2013), and orthographic typicality, describing the frequency and familiarity of certain word subcomponents being joined together (Woollams et al., 2011). |
| Picture Vocabulary Test (Gershon et al., 2013) | Language | Children hear a single spoken word presented with 4 images of objects, concepts, or actions, from which they must select the image most suited to the word. |
| Pattern Comparison Processing Speed Test (Carlozzi et al., 2015) | Processing Speed | Children must report whether two visual patterns are the same or not, varying in terms of colour, manipulation of object components, and quantity. |

**Table S2. Summary for Principal Component Analysis (PCA) of 7 NIH-TB Tests**

|  |  | Principal Component (PC) | | | | |
| --- | --- | --- | --- | --- | --- | --- |
| Cognitive Test Loading | **Cognitive Domain** | **PC1** | **PC2** | **PC3** | **PC4** | **PC5** |
| Picture Vocabulary | Language | 15.52 | 18.29 | 8.94 | 2.79 | 2.44 |
| Flanker | Attention | 14.16 | 15.95 | 2.94 | 39.29 | 3.80 |
| List Sorting | Working Memory | 16.55 | 7.29 | 4.97 | 9.01 | 61.17 |
| Dimensional Card Sorting | Set-Shifting | 16.57 | 15.77 | .24 | .36 | 2.79 |
| Pattern Comparison | Processing Speed | 11.49 | 24.07 | .39 | 45.56 | 9.10 |
| Picture Sequence Memory | Language | 10.64 | 1.38 | 69.59 | 1.31 | 15.36 |
| Oral Reading Recognition | Language | 15.06 | 17.24 | 12.95 | 1.70 | 5.35 |
| Variance Explained by PC (%) |  | 37.76 | 17.25 | 12.15 | 9.30 | 8.59 |
| Eigenvalue |  | 2.64 | 1.21 | .85 | .65 | .60 |

Note. “NIH-TB” = National Institutes of Health Toolbox

**Table S3. Polygenic score model specification for European and Non-European subsets.** For each model, 6 principal components (PC) of ancestry were included as covariates.

*Note.* ***** and ** represent significance at *p* < .01 and *p* < .001, respectively. Clumping thresholds of *p* = .1 and *p* = .2 were used for European and Non-European participants, respectively.

| Ancestry | Parameter | Coefficient | Standard Error | t-value | Pr(>\|t\|) |
| --- | --- | --- | --- | --- | --- |
| European | Intercept | .456 | 2.924e^-2^ | 15.589 | 1.307e^-53^ ****** |
|  | *g* Factor Loading | 26234.927 | 1824.743 | 14.377 | 5.196e^-46^ ****** |
|  | Sex (Male) | -1.011e^-1^ | 4.029e^-2^ | -2.510 | .012 ***** |
|  | PC1 | 3.008 | 1.526 | 1.971 | .049 |
|  | PC2 | -14.120 | 1.470 | -9.601 | 1.183e^-21^ ****** |
|  | PC3 | 5.241 | 1.600 | 3.276 | .001 ****** |
|  | PC4 | -1.685 | 1.469 | -1.146 | .252 |
|  | PC5 | -3.393 | 1.472 | -2.305 | .021 |
|  | PC6 | -6.737 | 1.476 | -4.564 | 5.130e^-6^ ****** |
| Non-European | Intercept | -.372 | 6.167e^-2^ | -6.026 | 2.235e^-9^ ****** |
|  | *g* Factor Loading | 18998.573 | 4164.188 | 4.564 | 5.578e^-6^ ****** |
|  | Sex (Male) | -.121 | .008 | -1.432 | .152 |
|  | PC1 | .980 | 1.505 | .652 | .515 |
|  | PC2 | 4.892 | 1.494 | 3.274 | .001 ****** |
|  | PC3 | -7.633 | 1.454 | -5.249 | 1.810e^-7^ ****** |
|  | PC4 | -2.006 | 1.461 | -1.373 | .170 |
|  | PC5 | -2.519 | 1.475 | -1.708 | .088 |
|  | PC6 | -.539 | 1.454 | -.371 | .711 |

**Table S4. Polygenic model performance for varying clumping thresholds for European and Non-European subsets.**

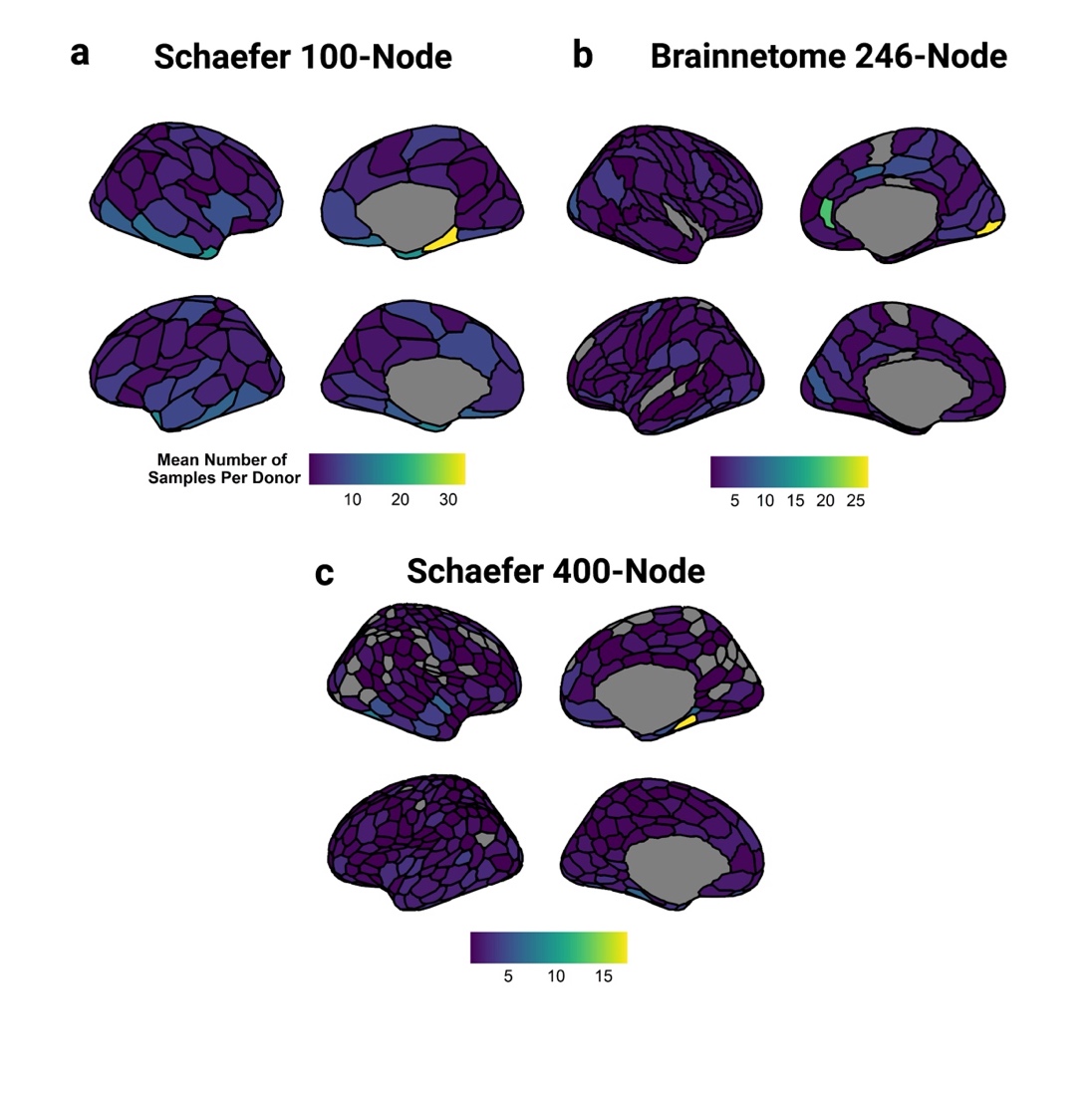

| Ancestry | Polygenic Clumping Threshold P_T_ | Polygenic Model Fit *R^2^* | *p* | *β* | Standard Error |
| --- | --- | --- | --- | --- | --- |
| European | .001 | .025 | 3.359e^-32^ | 2225.124 | 187.124 |
|  | .05 | .038 | 1.969e^-48^ | 9579.576 | 648.438 |
|  | .1 | .040 | 5.070e^-51^ | 13024.307 | 857.635 |
|  | .2 | .039 | 7.729e^-50^ | 17479.276 | 1165.373 |
|  | .3 | .038 | 6.215e^-48^ | 20755.122 | 1412.636 |
|  | .4 | .037 | 2.958e^-47^ | 23757.929 | 1629.252 |
|  | .5 | .036 | 5.196e^-46^ | 26234.927 | 1824.743 |
| Non-European | .001 | .012 | 1.053e^-4^ | 1637.998 | 420.961 |
|  | .05 | .012 | 1.188e^-4^ | 5410.462 | 1401.212 |
|  | .1 | .014 | 3.557e^-5^ | 7804.361 | 1880.474 |
|  | .2 | .017 | 2.867e^-6^ | 12194.429 | 2593.133 |
|  | .3 | .017 | 4.226e^-6^ | 14646.042 | 3169.217 |
|  | .4 | .016 | 9.407e^-6^ | 16434.281 | 3693.490 |
|  | .5 | .016 | 5.578e^-6^ | 18998.574 | 4164.188 |

**Figure S1. Distribution of mean number of samples per donor in the Allen Human Brain Atlas (AHBA) for 3 parcellations of increasing granularity.** (a) The Schaefer 100-node parcellation (Schaefer et al., 2018) has full AHBA spatial coverage, whilst this coverage decreases for the (b) Brainnetome 246-node (Fan et al., 2016) and (c) Schaefer 400-node parcellations (Schaefer et al., 2018), respectively. See Hawrylycz and colleagues (2012) for technical AHBA details.

**Table S5. Descriptive statistics for the distributions of local and global graph theory measures across 3 parcellations of varying granularity.** All are local measures, except density, and are defined in the Results. Note that the modularity statistic provided is a nodal measure of optimal community structure.

|  | Parcellation | | | | | |
| --- | --- | --- | --- | --- | --- | --- |
|  | Schaefer 100-Node | | Brainnetome 246-Node | | Schaefer 400-Node | |
| Metric | Mean (SD) | Range | Mean (SD) | Range | Mean (SD) | Range |
| Density | 6.55%  (.48) | 2.24 - 7.35 | 2.04% (.19) | .34 - 2.35 | .49% (.05) | .03 - .59 |
| Degree | 6.49 (3.17) | .65 - 15.57 | 4.99 (3.38) | 0 - 19.91 | 1.95 (2.08) | 0 - 11.30 |
| Clustering | .42 (.18) | 0 - .89 | .34 (.20) | 0 - .78 | .15 (.18) | 0 - .76 |
| Betweenness Centrality | 239.44 (290.42) | 0 - 1583.39 | 824.24 (996.12) | 0 - 6494.36 | 390.59 (812.89) | 0 -6910.37 |
| Eigenvector Centrality | .07 (.07) | 0 - .24 | .03 (.05) | 0 - .31 | .01 (.04) | 0 - .23 |
| Modularity | 3.47 (.33) | 2.89 - 4.01 | 5.76 (.53) | 3.80 - 6.88 | 10.28 (3.29) | 4.81 - 15.71 |
| Local Efficiency | .57 (.21) | 0 - .91 | .43 (.24) | 0 - .84 | .17 (.22) | 0 - .82 |
| Mean Total  Edge Length | 246.10 (177.23) | 11.42 - 1077.48 | 163.01 (140.50) | 0 - 736.61 | 44.84 (50.97) | 0 - 257.05 |

**Figure S2. Seed network for the Schaefer 100-node parcellation.** This seed network was generated by first thresholding individual connectomes in three steps: a 60% consensus threshold to remove spurious connections, thresholding at 27 streamlines, and then a 95% consensus threshold. This produced a seed network with 1.576% density, corresponding to 157 connections. Clusters of connections consistent across participants included the left auditory and secondary somatosensory divisions of the somatomotor cortex, namely the primary somatomotor network, alongside the right extra-striatal cortices and striate calcarine, namely the primary salience and attention network.

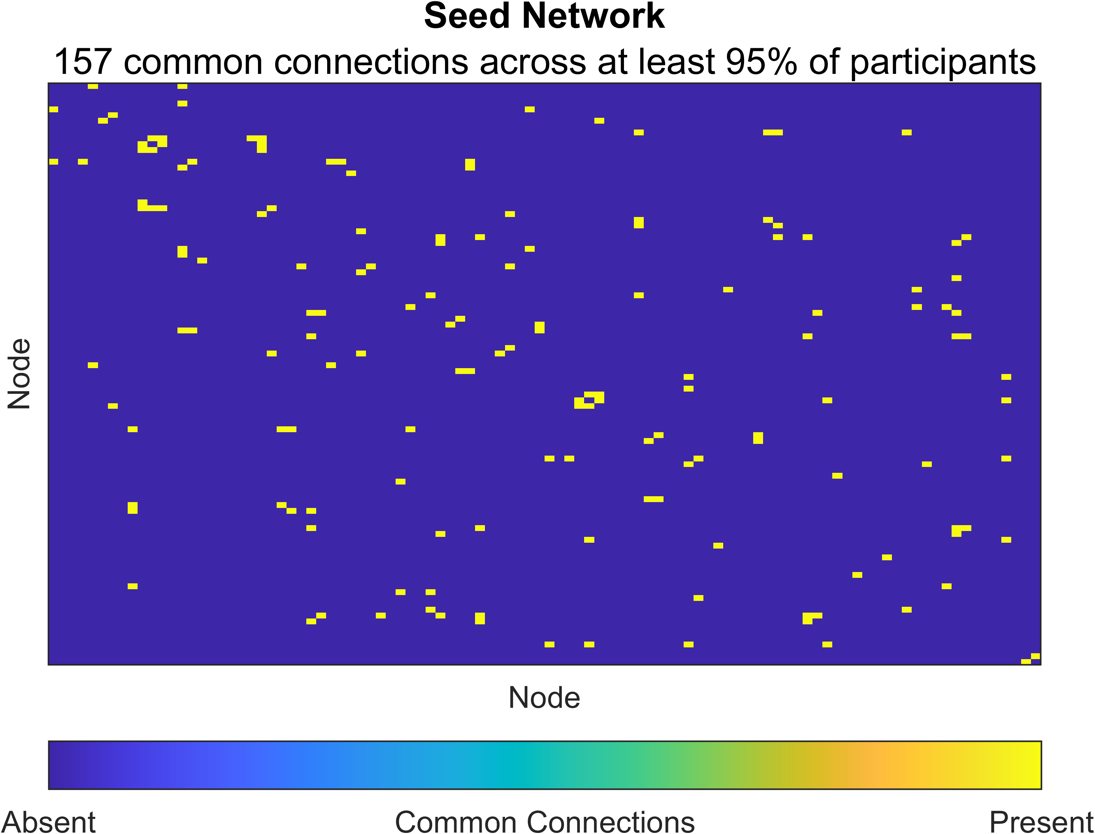

**Table S6. Lowest-energy group simulations with associated optimal η and γ parameters, across 3 parcellations.** Each generative rule was evaluated for 99,856 unique combinations of η [-7 ≤ η ≤ 7] and γ [-7 ≤ γ ≤ 7].

|  | Parcellation | | | | | | | | |
| --- | --- | --- | --- | --- | --- | --- | --- | --- | --- |
|  | Schaefer 100-Node | | | Brainnetome 246-Node | | | Schaefer 400-Node | | |
| Rule | Energy | η | γ | Energy | η | γ | Energy | η | γ |
| Clu-Avg | .120 | -6.422 | -4.822 | .228 | -4.200 | -2.289 | .173 | -5.089 | 1.000 |
| Deg-Avg | .148 | -5.578 | 1.667 | .240 | -4.556 | 1.978 | .138 | -3.533 | 1.444 |
| Matching | .100 | -3.800 | .333 | .106 | -2.467 | .333 | .110 | -2.956 | .467 |
| Neighbours | .090 | -2.911 | .244 | .110 | -2.200 | .289 | .107 | -2.911 | .378 |
| Spatial | .205 | -6.378 | 4.289 | .289 | -4.111 | .244 | .268 | -3.578 | 3.400 |

*Note.* “Clu-Avg” = Average Clustering Coefficient, “Deg-Avg” = Average Degree.

**Table S7. Mean topological dissimilarity (TD) and correlations between simulated and observed degree for 1000 simulations of each generative model’s lowest energy η and γ combination, across 3 parcellations.** For the Brainnetome 246-node parcellation, the models differed significantly in their topological dissimilarity [*F*(4,4999) = 1654.00, *p* < .001] and ability to capture observed degree [*F*(4,4999) = 1643.50, *p* < .001], with all post-hoc comparisons highly significant (*p* < .001). For the Schaefer 400-node parcellation, the models also differed significantly in their topological dissimilarity [*F*(4,4999) = 4502.14, *p* < .001], with all post-hoc comparisons highly significant (*p* < .001). The models also differed in their correlation with observed degree [*F*(4,4999) = 2197.93, *p* < .001], with all post-hoc comparisons highly significant, apart from the two homophily models performing similarly to each other (*p* = .991).

|  | Schaefer 100-Node | |
| --- | --- | --- |
| GNM Rule | Topological Dissimilarity  (Mean ± SD) | Simulated – Observed Degree Pearson’s *r* (Mean ± SD) |
| Clu-Avg | 1.975 ± .156 | .165 ± .071 |
| Deg-Avg | .963 ± .236 | .343 ± .062 |
| Matching | 1.004 ± .280 | .394 ± .079 |
| Neighbours | .979 ± .265 | .412 ± .077 |
| Spatial | 1.058 ± .339 | .130 ± .062 |
|  | **Brainnetome 246-Node** | |
| Clu-Avg | 1.683 ± .132 | .050 ± .062 |
| Deg-Avg | 1.462 ± .264 | .076 ± .031 |
| Matching | 1.035 ± .239 | .119 ± .046 |
| Neighbours | 1.098 ± .252 | .129 ± .050 |
| Spatial | .973 ± .281 | -.018 ± .039 |
|  | **Schaefer 400-Node** | |
| Clu-Avg | 1.482 ± .188 | -.095 ± .120 |
| Deg-Avg | .998 ± .160 | .197 ± .044 |
| Matching | .618 ± .187 | .016 ± .070 |
| Neighbours | .577 ± .194 | .015 ± .073 |
| Spatial | .763 ± .138 | -.055 ± .044 |

*Note.* “Clu-Avg” = Average Clustering Coefficient, “Deg-Avg” = Average Degree.

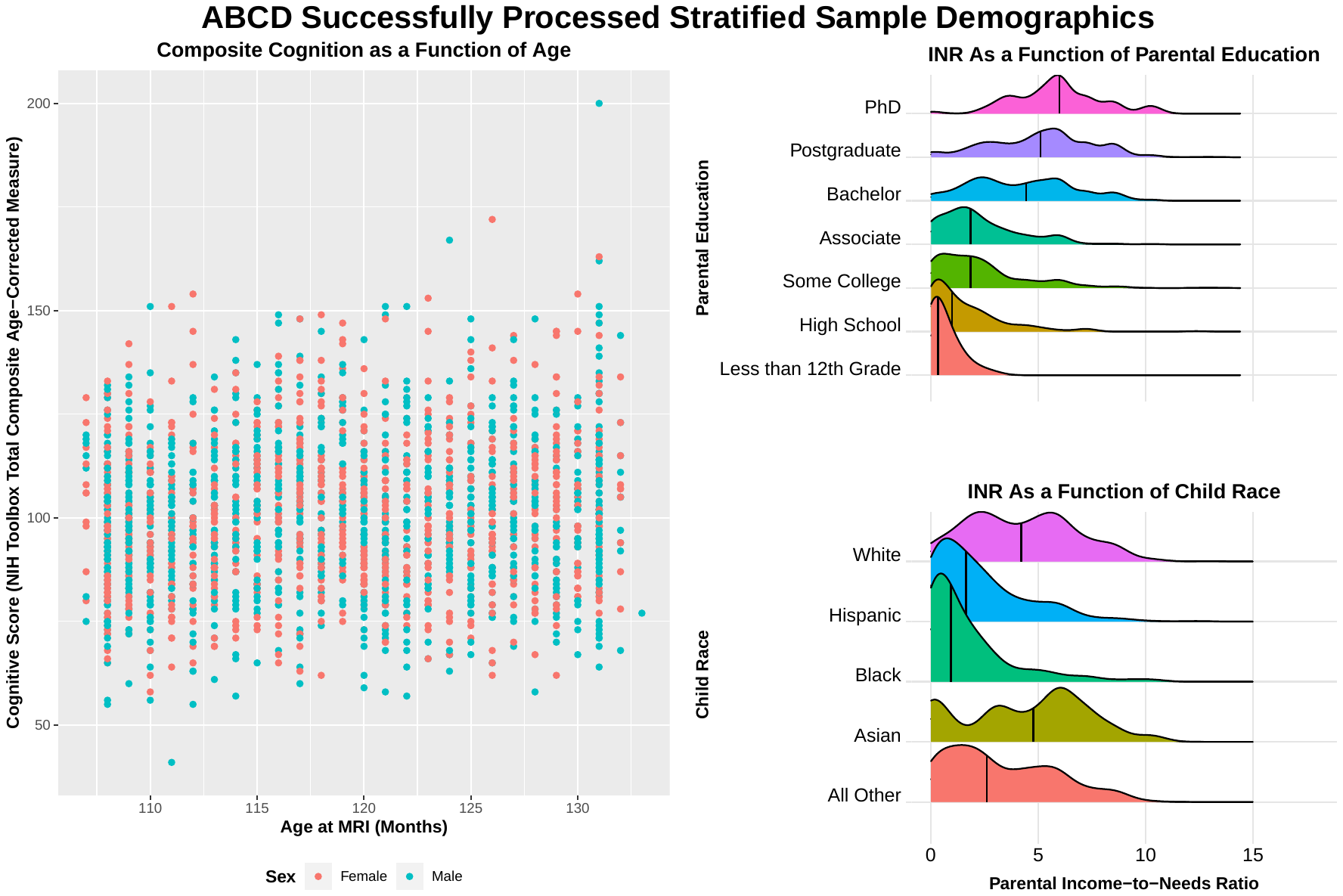
**Figure S3. Summary of stratified participant demographics.** **(a)** Relationship between NIH-total composite cognition scores and age scan, by sex. **(b)** and **(c)** show income-to-needs ratio (INR), an index of socio-economic status, as a function of parental education and child race, respectively. Vertical black lines within each distribution represent the median.

**b**

**a**

**c**

*Note.* “NIH” = National Institutes of Health; “MRI” = Magnetic resonance imaging.

**Table S8. Summary of Gene Ontologies for Parameterised Nodal Wiring Costs, Value, and Cognitive Ability.** Allen Human Brain Atlas (AHBA) genes predicted parameterised nodal wiring costs and values, separately, for each participant, through partial least squares regression. Following 10,000 permutations for each participant, AHBA genes with permuted *p*-values less than .05 across all participants were selected and then ranked by decreasing mean loading onto the first latent variable. 76,745 short-nucleotide polymorphisms were ranked by decreasing absolute β in the cognitive ability polygenic score. All gene lists were submitted separately to g:Profiler (Kolberg et al., 2020) for gene enrichment, with a cut-off of *p* < .05 corrected for multiple comparisons, and default parameters. Electronic annotations were excluded for robustness.

|  | Parameterised Nodal Wiring Costs | Parameterised Nodal Wiring Value | Cognitive Ability |
| --- | --- | --- | --- |
| Number of Genes | 951 | 561 | 15,234 |
| g:Profiler Link | https://biit.cs.ut.ee/gplink/l/m2E_uLK0TG | https://biit.cs.ut.ee/gplink/l/T8OtQThDQ_ | https://biit.cs.ut.ee/gplink/l/lGxCWYZpSo |
| % BP/CC/MF | 10.526% / 15.790% / 73.684% | 43.902% / 29.268% / 26.829% | 49.495% / 27.609% / 22.896% |
| Top 10 Enriched Categories | GO:0046873, Metal Ion Transmembrane Transporter Activity (*p_adj_* = 2.116e^-4^) | GO:0099536, Synaptic Signaling (*p_adj_* = 1.130e^-5^) | GO:0097060, Synaptic Membrane (*p_adj_* = 3.485e^-29^) |
|  | GO:0005215, Transporter Activity (*p_adj_* = 4.480e^-4^) | GO:0007267, Cell-Cell Signaling (*p_adj_* = 1.949e^-5^) | GO:0045202, Synapse (*p_adj_* = 2.586e^-26^) |
|  | GO:0008324, Cation Transmembrane Transporter Activity (*p_adj_* = 8.192e^-4^) | GO:0098916, Anterograde Trans-Synaptic Signaling (*p_adj_* = 2.348e^-5^) | GO:0098794, post-Synapse (*p_adj_* = 3.684e^-26^) |
|  | GO:0022890, Inorganic Cation Transmembrane Transporter Activity (*p_adj_* = 9.618e^-4^) | GO:0007268, Chemical Synaptic Transmission (*p_adj_* = 2.348e^-5^) | GO:0045211, Post-Synapse Membrane (*p_adj_* = 2.072e^-24^) |
|  | GO:0022857, Transmembrane Transporter Activity (*p_adj_* = .002) | GO:0099537, Trans-Synaptic Signaling (*p_adj_* = 3.284e^-5^) | GO:0043005, Neuron Projection (*p_adj_* = 1.492e^-20^) |
|  | GO:0015075, Ion Transmembrane Transporter Activity (*p_adj_* = .003) | GO:0071944, Cell Periphery (*p_adj_* = 1.169e^-4^) | GO:0099537, Trans-Synaptic Signaling (*p_adj_* = 3.821e^-20^) |
|  | GO:0015318, Inorganic Molecular Entity Transmembrane Transporter Activity (*p_adj_* = .004) | GO:0005886, Plasma Membrane (*p_adj_* = 4.230e^-4^) | GO:009536, Synaptic Signaling (*p_adj_* = 7.602e^-20^) |
|  | GO:008514, Organic Anion Transmembrane Transporter Activity (*p_adj_* = .005) | GO:0045202, Synapse (*p_adj_* = 5.586e^-4^) | GO:0098916, Anterograde Trans-Synaptic Signaling (*p_adj_* = 2.212e^-19^) |
|  | GO:0000149, SNARE Binding (*p_adj_* = .005) | GO:0043005, Neuron Projection (*p_adj_* = .002) | GO:0007268, Chemical Synaptic Transmission (*p_adj_* = 2.212e^-19^) |
|  | GO:0015291, Secondary Active Transmembrane Transporter Activity (*p_adj_* = .005) | GO:0022824, Transmitter-Gated Ion Channel Activity (*p_adj_* = .004) | GO:0030054, Cell Junction (*p_adj_* = 7.227e^-19^) |

*Note.* “MF” = Molecular function; “BP” = Biological processes; “CC” = Cellular Components.

**Table S9. Gene ontologies for a comparative pathway enrichment analysis between parameterised nodal wiring costs, value, and cognitive ability PGS.** We created 3 ranked gene lists of genes predictive of parameterised nodal wiring costs, value, and SNPs predictive of the cognitive ability PGS (see Table S9). These gene lists were inputted into a multi-query comparative pathway analysis in g:Profiler with default parameters, and a cut-off of *p* < .01. Electronic annotations were excluded.

|  | | *p* values | | |
| --- | --- | --- | --- | --- |
| GO ID | **Term Name** | Cost | Value | PGS |
| GO:0043005  CC | neuron projection | 1.000 | 0.002 | 1.573e^-10^ |
| GO:0030594  MF | neurotransmitter receptor activity | 1.000 | 0.013 | 1.846e^-9^ |
| GO:0034702  CC | ion channel complex | 1.000 | 0.173 | 5.946e^-9^ |
| GO:0099537  BP | trans-synaptic signaling | 1.000 | 3.284e^-5^ | 7.604e^-9^ |
| GO:0099536  BP | synaptic signaling | 1.000 | 1.130e^-5^ | 8.678e^-9^ |
| GO:0045202  CC | synapse | 0.324 | 0.001 | 1.301e^-8^ |
| GO:0034703  CC | cation channel complex | 1.000 | 0.168 | 1.330e^-8^ |
| GO:0098797  CC | plasma membrane protein complex | 1.000 | 0.456 | 1.735e^-8^ |
| GO:0007268  BP | chemical synaptic transmission | 1.000 | 2.348e^-5^ | 1.917e^-8^ |
| GO:0098916  BP | anterograde trans-synaptic signaling | 1.000 | 2.348e^-5^ | 1.917e^-8^ |
| GO:0007215  BP | glutamate receptor signaling pathway | 1.000 | 1.000 | 1.020e^-7^ |
| GO:0098794  CC | post synapse | 1.000 | 0.403 | 1.620e^-7^ |
| GO:0022836  MF | gated channel activity | 1.000 | 0.056 | 1.783e^-7^ |
| GO:0005230  MF | extracellular ligand-gated ion channel activity | 1.000 | 0.014 | 2.613e^-7^ |
| GO:0005272  MF | sodium channel activity | 1.000 | 1.000 | 5.191e^-7^ |
| GO:0022835  MF | transmitter-gated channel activity | 1.000 | 0.004 | 8.765e^-7^ |
| GO:0022824  MF | transmitter-gated ion channel activity | 1.000 | 0.004 | 8.765e^-7^ |
| GO:0035249  BP | synaptic transmission, glutamatergic | 1.000 | 1.000 | 1.466e^-6^ |
| GO:0045211  CC | postsynaptic membrane | 1.000 | 1.000 | 1.705e^-6^ |
| GO:0034706  CC | sodium channel complex | 1.000 | 0.271 | 1.851e^-6^ |
| GO:0098988  MF | G protein-coupled glutamate receptor activity | 1.000 | 1.000 | 4.098e^-6^ |
| GO:0001640  MF | adenylate cyclase inhibiting G protein-coupled glutamate receptor activity | 1.000 | 1.000 | 4.098e^-6^ |
| GO:0015081  MF | sodium ion transmembrane transporter activity | 0.315 | 1.000 | 4.676e^-6^ |
| GO:0005216  MF | ion channel activity | 1.000 | 0.032 | 4.712e^-6^ |
| GO:0005261  MF | cation channel activity | 1.000 | 1.000 | 9.966e^-6^ |
| GO:0097060  CC | synaptic membrane | 1.000 | 1.000 | 1.042e^-5^ |
| GO:0022834  MF | ligand-gated channel activity | 1.000 | 0.006 | 1.052e^-5^ |
| GO:0015276  MF | ligand-gated ion channel activity | 1.000 | 0.006 | 1.052e^-5^ |
| GO:0030424  CC | axon | 0.019 | 0.473 | 1.215e^-5^ |
| GO:0046873  MF | metal ion transmembrane transporter activity | 2.116e^-4^ | 1.000 | 1.234e^-5^ |
| GO:0001518  CC | voltage-gated sodium channel complex | 1.000 | 0.378 | 1.755e^-5^ |
| GO:0007267  BP | cell-cell signaling | 1.000 | 1.949e^-5^ | 0.270 |
| GO:0042995  CC | cell projection | 1.000 | 0.010 | 2.142e^-5^ |
| GO:0120025  CC | plasma membrane bounded cell projection | 1.000 | 0.006 | 2.563e^-5^ |
| GO:0042391  BP | regulation of membrane potential | 1.000 | 1.000 | 3.514e^-5^ |
| GO:0015267  MF | channel activity | 1.000 | 0.063 | 4.425e^-5^ |
| GO:0004970  MF | ionotropic glutamate receptor activity | 1.000 | 1.000 | 4.451e^-5^ |
| GO:0036477  CC | Somato-dendritic compartment | 1.000 | 0.138 | 4.665e^-5^ |
| GO:0022803  MF | passive transmembrane transporter activity | 1.000 | 0.064 | 4.815e^-5^ |
| GO:0051966  BP | regulation of synaptic transmission, glutamatergic | 1.000 | 1.000 | 4.958e^-5^ |
| GO:0086010  BP | membrane depolarization during action potential | 1.000 | 1.000 | 1.019e^-4^ |
| GO:0097447  CC | dendritic tree | 1.000 | 0.092 | 1.121e^-4^ |
| GO:0071944  CC | cell periphery | 1.000 | 1.169e^-4^ | 0.541 |
| GO:0098960  MF | postsynaptic neurotransmitter receptor activity | 1.000 | 0.907 | 1.448e^-4^ |
| GO:0035725  BP | sodium ion transmembrane transport | 1.000 | 1.000 | 1.534e^-4^ |
| GO:0005248  MF | voltage-gated sodium channel activity | 1.000 | 1.000 | 1.620e^-4^ |
| GO:0050804  BP | modulation of chemical synaptic transmission | 1.000 | 1.000 | 1.757e^-4^ |
| GO:0030425  CC | dendrite | 1.000 | 0.082 | 1.918e^-4^ |
| GO:0099177  BP | regulation of trans-synaptic signaling | 1.000 | 1.000 | 1.992e^-4^ |
| GO:1902495  CC | transmembrane transporter complex | 0.264 | 1.000 | 2.214e^-4^ |
| GO:0007216  BP | G protein-coupled glutamate receptor signaling pathway | 1.000 | 1.000 | 3.482e^-4^ |
| GO:0005886  CC | plasma membrane | 1.000 | 4.230e^-4^ | 1.000 |
| GO:0005215  MF | transporter activity | 0.001 | 1.000 | 1.000 |
| GO:0006814  BP | sodium ion transport | 1.000 | 0.844 | 0.001 |
| GO:0008328  CC | ionotropic glutamate receptor complex | 1.000 | 0.079 | 0.001 |
| GO:0015318  MF | inorganic molecular entity transmembrane transporter activity | 0.004 | 0.176 | 0.001 |
| GO:0022843  MF | voltage-gated cation channel activity | 1.000 | 1.000 | 0.001 |
| GO:0022890  MF | inorganic cation transmembrane transporter activity | 0.001 | 1.000 | 0.001 |
| GO:0098878  CC | neurotransmitter receptor complex | 1.000 | 0.180 | 0.001 |
| GO:0008324  MF | cation transmembrane transporter activity | 0.001 | 1.000 | 0.006 |
| GO:1990351  CC | transporter complex | 0.252 | 1.000 | 0.001 |
| GO:0001505  BP | regulation of neurotransmitter levels | 0.304 | 1.000 | 0.002 |
| GO:0019228  BP | neuronal action potential | 1.000 | 1.000 | 0.002 |
| GO:0008038  BP | neuron recognition | 1.000 | 1.000 | 0.002 |
| GO:0022857  MF | transmembrane transporter activity | 0.002 | 1.000 | 0.643 |
| GO:0015075  MF | ion transmembrane transporter activity | 0.003 | 0.513 | 0.027 |
| GO:0007156  BP | homophilic cell adhesion via plasma membrane adhesion molecules | 1.000 | 1.000 | 0.003 |
| GO:0005343  MF | organic acid: sodium symporter activity | 1.000 | 1.000 | 0.003 |
| GO:0030054  CC | cell junction | 1.000 | 0.005 | 0.004 |
| GO:0005244  MF | voltage-gated ion channel activity | 1.000 | 1.000 | 0.004 |
| GO:0022832  MF | voltage-gated channel activity | 1.000 | 1.000 | 0.004 |
| GO:0005283  MF | amino acid: sodium symporter activity | 1.000 | 1.000 | 0.004 |
| GO:0008514  MF | organic anion transmembrane transporter activity | 0.005 | 1.000 | 1.000 |
| GO:0015370  MF | solute: sodium symporter activity | 0.168 | 1.000 | 0.005 |
| GO:0000149  MF | SNARE binding | 0.005 | 1.000 | 1.000 |
| GO:0001504  BP | neurotransmitter uptake | 1.000 | 1.000 | 0.005 |
| GO:0031226  CC | intrinsic component of plasma membrane | 0.979 | 0.226 | 0.005 |
| GO:0008509  MF | anion transmembrane transporter activity | 0.005 | 1.000 | 1.000 |
| GO:0007196  BP | adenylate cyclase-inhibiting G protein-coupled glutamate receptor signaling pathway | 1.000 | 1.000 | 0.006 |
| GO:0004930  MF | G protein-coupled receptor activity | 1.000 | 1.000 | 0.006 |
| GO:0015291  MF | secondary active transmembrane transporter activity | 0.006 | 1.000 | 0.238 |
| GO:0005887  CC | integral component of plasma membrane | 0.699 | 0.429 | 0.006 |
| GO:0015277  MF | kainate selective glutamate receptor activity | 1.000 | 1.000 | 0.007 |
| GO:0098793  CC | Pre-synapse | 0.007 | 0.028 | 0.118 |
| GO:0044304  CC | main axon | 1.000 | 1.000 | 0.007 |
| GO:0003231  BP | cardiac ventricle development | 1.000 | 0.008 | 1.000 |
| GO:0099094  MF | ligand-gated cation channel activity | 1.000 | 1.000 | 0.008 |
| GO:0071870  BP | cellular response to catecholamine stimulus | 1.000 | 0.253 | 0.008 |
| GO:0071869  BP | response to catecholamine | 1.000 | 0.253 | 0.008 |
| GO:0071868  BP | cellular response to monoamine stimulus | 1.000 | 0.253 | 0.008 |
| GO:0071867  BP | response to monoamine | 1.000 | 0.253 | 0.008 |
| GO:0048699  BP | generation of neurons | 1.000 | 0.017 | 0.008 |
| GO:0099095  MF | ligand-gated anion channel activity | 1.000 | 0.009 | 0.086 |
| GO:1904315  MF | transmitter-gated ion channel activity involved in regulation of postsynaptic membrane potential | 1.000 | 1.000 | 0.009 |
| GO:0099529  MF | neurotransmitter receptor activity involved in regulation of postsynaptic membrane potential | 1.000 | 1.000 | 0.009 |

*Note.* “MF” = Molecular function; “BP” = Biological processes; “CC” = Cellular Components; “GO” = Gene Ontology; “PGS” = Polygenic score.
